# Flattening of diurnal glucocorticoid oscillations causes Cd36 and insulin-mediated obesity

**DOI:** 10.1101/2020.01.02.893081

**Authors:** Stefan Tholen, Kyle M. Kovary, Atefeh Rabiee, Ewa Bielczyk-Maczyńska, Wenting Yang, Fredric B. Kraemer, Mary N. Teruel

## Abstract

Chronic stressors flatten diurnal glucocorticoid oscillations, resulting in obesity, insulin resistance, and diabetes. How such flattened diurnal glucocorticoid oscillations increase fat storage is unknown. Here we investigated the consequences in mice and found that flattening of glucocorticoid oscillations results not only in body weight gain, mainly due to increases in white fat depot mass, but also leads to hyperinsulinemia and fat accumulation in brown adipose tissue. A transcriptomic analysis of white and brown adipose tissues revealed that flattened glucocorticoid oscillations cause dysregulated lipid metabolism with a prominent role of the fatty acid transporter Cd36 and insulin-driven adipocyte hypertrophy. Indeed, *Cd36* knockout mice are partially protected against the adverse effects of flattened GC oscillations including body weight gain and lipid accumulation in the brown and visceral white fat depots. These results show the molecular mechanisms how flattened glucocorticoid oscillations can cause obesity and diabetes.

## INTRODUCTION

Glucocorticoids are steroid hormones that are capable of causing tremendous shifts in carbohydrate and fatty acid metabolism throughout the body (Macfarlane et al., 2008). Glucocorticoids act on multiple tissues to increase available circulating energy substrates by promoting catabolism and deferring anabolism. A fundamental aspect of glucocorticoid regulation is that glucocorticoids are not maintained at constant levels in healthy mammals, but instead are secreted in diurnal oscillations (Weitzman et al., 1971) (Figure 1A, grey pattern). The diurnal pattern is essential for normal body function: the daily increase in glucocorticoids provides a wake-up signal and turns on whole body metabolism and the daily trough, or off-period, allows a rest period in which catabolic glucocorticoid-driven processes are not activated and the body can recover and reset (Dallman et al., 2000; Oster et al., 2017; Sapolsky et al., 1986).

**Figure 1.**
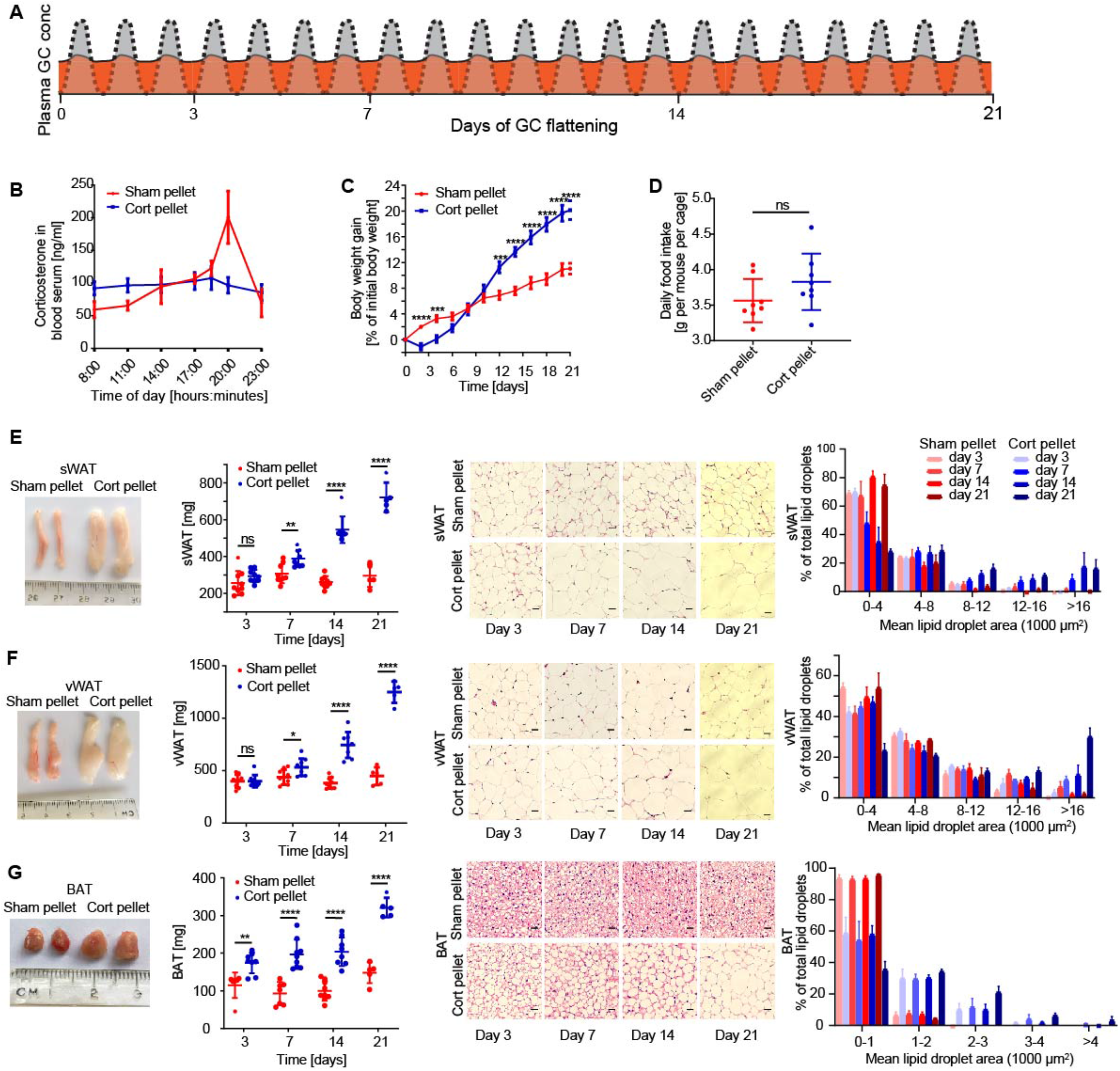
Flattening of circadian glucocorticoid oscillations causes rapid adipocyte hypertrophy. **(A)** Schematic of approach used to determine how flattening of glucocorticoid oscillations affects adipocyte hypertrophy over time. The normal circadian GC oscillation pattern is shown in grey. The flattened glucocorticoid pattern is shown in orange. **(B)** Time course of average corticosterone levels in mice implanted with corticosterone pellets (cort-pellet mice) compared to mice implanted with sham pellets (sham-pellet mice) 7 days after pellet implantation. n = 8 for sham pellet; n = 8 for corticosterone pellet, mean ± SEM. **(C)** Time course showing body weight gain of cort-pellet compared to sham-pellet mice. n = 11, mean ± SEM. Unpaired t test (sham pellet versus cort pellet), *p < 0.05, **p < 0.01, ***p < 0.001, ****p < 0.0001. **(D)** Average daily food intake of each mouse per cage (n = 8). Up to 5 mice were housed in each cage. Food intake was monitored for 21 days, mean ± SD. Unpaired t-test, ns = not significant. **(E-G)** sWAT, vWAT, and BAT fat pad images, fat pad weight, and hematoxylin and eosin (H & E) staining analysis. Fat pad images were taken 21 days after pellet implantation. Fat pad weight was measured at 3, 7, 14, and 21 days after pellet implantation, n = 5-9, mean ± SD. Unpaired t test, **p < 0.01, ****p < 0.0001. One representative H & E staining is shown for each timepoint (n = 3). Scale bars 50 µm. See also Table S1.

The diurnal glucocorticoid pattern is controlled by the hypothalamic-pituitary-adrenal (HPA) axis which secretes glucocorticoids in response to light-dark signals from the master clock in the brain (Spiga et al., 2014). This same HPA axis also secretes glucocorticoids in response to stress. Short term, stress-mediated spikes in glucocorticoids are beneficial to increase overall catabolic activity and ready the body for a fight-or-flight response (Sapolsky et al., 1986). However, if the stress becomes chronic, the continual secretion of glucocorticoids flattens the normal diurnal oscillatory pattern, meaning that peak levels are decreased and trough levels are elevated, without the daily mean values significantly changing (Dallman et al., 2000) (Figure 1A, orange pattern). Such flattening of glucocorticoid oscillations can occur from depression, jet lag, aging, and irregular eating and sleeping schedules (Adam, 2017; Leproult et al., 1997), as well as in disease conditions such as Cushing’s (Rossi et al., 2000), and has been shown to cause obesity and metabolic disease (Broussard and Van Cauter, 2016; Campbell et al., 2011; Dallman et al., 2000; Joseph and Golden, 2017; Mazziotti et al., 2011).

Fat mass expansion can occur both through adipogenesis, or the generation of new adipocytes from precursor cells, and from enlargement of existing adipocytes (hypertrophy)(Ghaben and Scherer, 2019). Previously we had studied the mechanisms controlling adipogenesis when glucocorticoids are flattened and found that the cause of the increased adipogenesis is that preadipocytes have a transcriptional network with slow and fast positive feedbacks centered on PPARG, the master regulator of adipogenesis, responses selectively to flattened, but not oscillating diurnal, glucocorticoid signals (Bahrami-Nejad et al., 2018). These results provided a molecular mechanism for the adipogenesis observed when glucocorticoid oscillations are flattened. However, the molecular mechanisms underlying the significant adipocyte hypertrophy when glucocorticoids are flattened remained poorly understood. Given the increasing evidence that the harmful metabolic effects of obesity originate primarily from adipocyte hypertrophy, rather than from adipogenesis (Ghaben and Scherer, 2019), it is of broad interest to determine the cause of the adipocyte hypertrophy when glucocorticoid oscillations are flattened.

That adipose tissue mass and adipocyte hypertrophy can increase when glucocorticoid oscillations are flattened (Bahrami-Nejad et al., 2018; Dallman et al., 2000; Karatsoreos et al., 2010; Rebuffé-Scrive et al., 1992; Shen et al., 2017) is puzzling since glucocorticoids are catabolic and generally drive breakdown of tissue mass. Indeed, other studies have shown that animals and humans can lose adipose tissue mass under some conditions when glucocorticoids are flattened (Lee et al., 2018; Peckett et al., 2011). We hypothesized that carrying out timecourse measurements would allow us to resolve these controversial results. In particular, we noted that the latter studies typically used high pharmacological doses of synthetic glucocorticoids or administered glucocorticoids together with a high fat diet which introduce confounding factors (Dunford and Riddell, 2016). Thus, to understand how obesity and adipocyte hypertrophy is generated just by glucocorticoid flattening, we used pellets containing corticosterone, the main physiological glucocorticoid in mice (Sapolsky et al., 1986), rather than synthetically stabilized types of glucocorticoids. We also avoided using high pharmacological doses and instead used experimental conditions that maintained normal mean circulating levels of glucocorticoids.

We designed timecourse experiments to mimic a limited period of chronic stress that flattens glucocorticoid levels and we sought to answer three main questions: (i) how long does it take after glucocorticoid flattening or glucocorticoid peak increases for adipocytes to become hypertrophic, (ii) how long after glucocorticoid flattening do markers for metabolic disease in the plasma and liver start to appear, and (iii) what is the mechanism that increases lipid storage in the hypertrophic adipocytes. We found that within 3 days of glucocorticoid flattening, mice develop hyperinsulinemia and strongly upregulate de novo fatty acid synthesis genes in white and brown adipose tissue as determined by RNAseq. This latter result is surprising since lipid accumulation in adipocytes is thought to be generated primarily from fatty acids taken up from the circulation (Santoro et al., 2021; Strawford et al., 2004). Acute increases in brown adipose tissue mass and adipocyte hypertrophy were observed starting at 3 days, followed by acute increases in white fat mass and adipocyte hypertrophy starting at 7 days. Despite a more than 2-fold increase in the brown and white fat mass size within 21 days, no significant increase in circulating fatty acids or accumulation of lipid in the liver was observed, supporting that the hyperinsulinemia and upregulation of de novo fatty acid synthesis in adipocytes drive the hypertrophy. Experiments in mice in which the fatty acid transporter Cd36 was knocked out support that increased fatty acid uptake also partially contributes to the adipocyte hypertrophy after glucocorticoid flattening. Surprisingly, the hyperinsulinemia and much of the adipocyte hypertrophy were reversed within 3 weeks after pellets stopped releasing glucocorticoid. Since the rapid increase in adipocyte size results from uptake of both fatty acid and glucose from the circulation, the hyperinsulinemia and adipocyte hypertrophy resulting from losing the glucocorticoid troughs likely have initially a protective role to keep circulating fatty acids and glucose low. In this way, the rapid increase in fat mass may serve to counteract the potentially harmful side effects of the catabolic activity increase resulting from temporary stress and continuous glucocorticoids, as long as the stress conditions do not exceed a few weeks.

## RESULTS

### Flattening of diurnal glucocorticoid oscillations causes rapid adipocyte hypertrophy in white and brown fat

To mimic the chronic stress condition, we flattened daily glucocorticoid oscillations in C57BL/6J male mice by implanting pellets that release corticosterone, the main glucocorticoid in mice, at a constant rate over 21 days. We specifically chose the dose of glucocorticoids that was released each day such that circulating glucocorticoid oscillations were flattened but the mean glucocorticoid level was kept approximately at normal physiological levels (Bahrami-Nejad et al., 2018; Hodes et al., 2012). As shown in Figure 1B, blood serum ELISA measurements confirmed that mice implanted with corticosterone pellets (hereafter referred to as “cort-pellet mice”) had decreased peak and increased trough levels of glucocorticoids compared to sham pellet-implanted mice (hereafter referred to as “sham-pellet mice”). Even though the mean glucocorticoid levels between the cort-pellet and sham-pellet mice were similar throughout the 21-day experiment, the mice with flattened glucocorticoid oscillations significantly increased their body weight compared to sham-pellet mice and ended up weighing 9.1% more (Figure 1C). As found in our previous study (Bahrami-Nejad et al., 2018) and in other studies which administered sustained glucocorticoids longterm (Bowles et al., 2015; Luijten et al., 2019), the increased weight gain in the cort-pellet mice cannot be explained by higher food intake (Figure 1d), and a reduction in metabolism may be a more significant contributing factor.

To assess whether the cort-pellet mice gain weight mainly by expansion of fat depots, we dissected and weighed the main white adipose tissue (WAT) and brown adipose tissue (BAT) depots in the mice. As the main subcutaneous WAT depot, we used the inguinal WAT which is a located in the flank region under the skin (Chusyd et al., 2016). As the main visceral WAT depot, we used the epididymal WAT which surrounds the gonads (Chusyd et al., 2016). We will hereafter refer to inguinal subcutaneous WAT as “sWAT”, and epididymal visceral WAT as “vWAT”. As the main brown fat depot, we used the interscapular brown fat depot which is the largest and most accessible brown fat depot in rodents (Cannon and Nedergaard, 2004). Strikingly, sWAT and vWAT weight increased more than two-fold and BAT weight increased three-fold within 21 days of glucocorticoid oscillation flattening (Figure 1E-1G, left two panels). Significant expansions of sWAT and vWAT were apparent after 7 days of glucocorticoid flattening, and an even earlier expansion was observed in BAT after only 3 days (Figure 1E-1G, left two panels). The expansions of WAT and BAT were confirmed by hematoxylin and eosin (H&E) staining which revealed significantly increased cell volumes of the adipocytes in the BAT from day 3 and in the WAT from day 7 (Figure 1E-1G, right two panels).

As a control to test whether the lipid accumulation in WAT and BAT shown in Figure 1 is due to flattening of diurnal glucocorticoid oscillations and not caused by the amount of corticosterone released into the mouse by the pellet, we injected C57BL/6J male mice at 5 PM each day for 21 days with the same total amount of corticosterone that is released by the pellet in one day (Figure 2A). This injection resulted in a more than 10-fold increase in the daily peak amplitude of corticosterone while still maintaining the trough period (Figure 2B). Markedly, despite the higher peak level, corticosterone-injected mice did not show a significant increase in body weight compared to sham-injected control mice (Figure 2C). Furthermore, their food intake was not altered (Figure 2D). In addition, no differences in fat pad weight were detected between the sWAT, vWAT, or BAT of the cort-and sham-injected mice (Figure 2E-G), and histological analysis of the tissues showed no differences in lipid accumulation (Figure S1).

**Figure 2.**
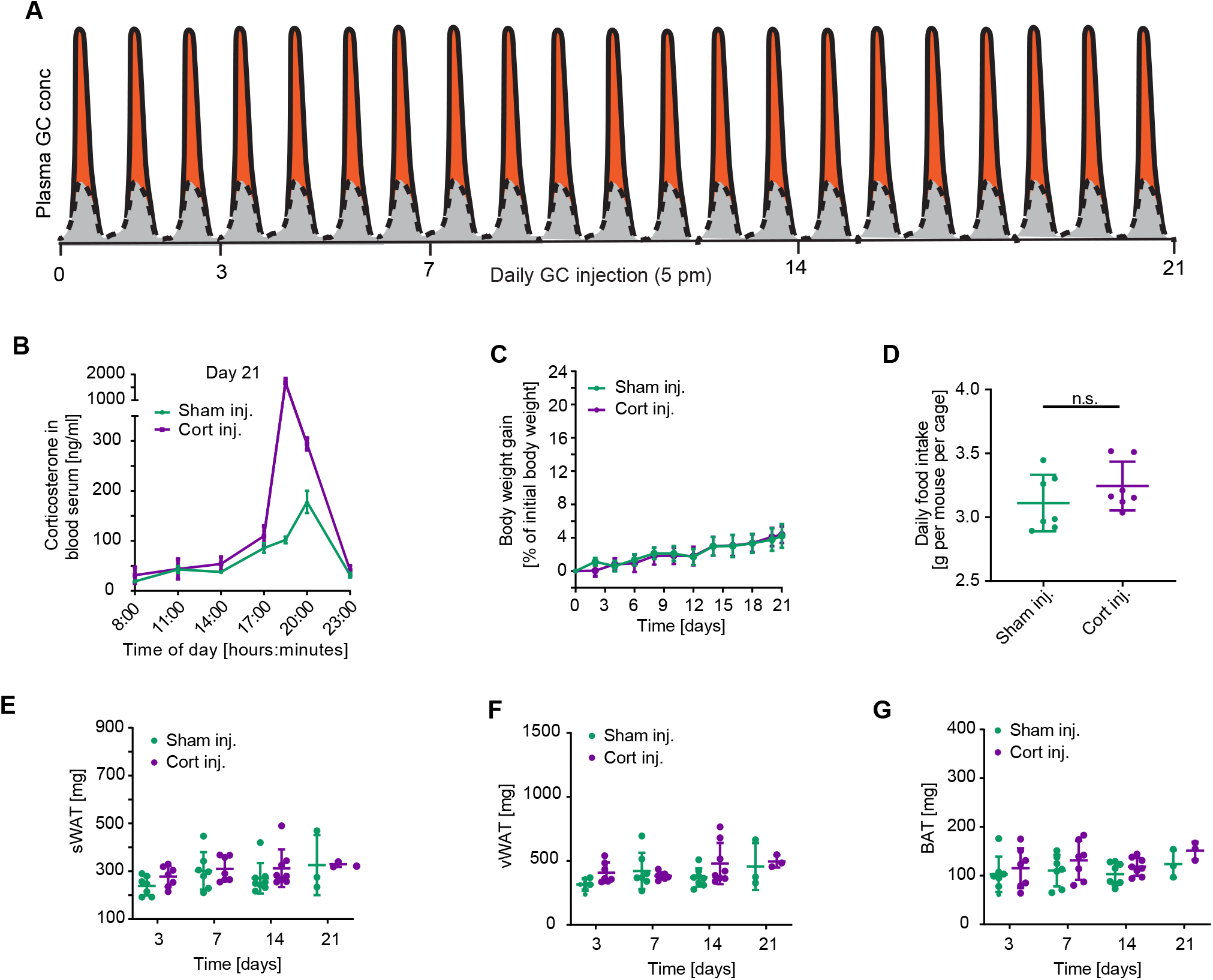
Increasing the peak amplitude of corticosterone in the evening phase does not affect body weight gain or lipid accumulation in WAT. **(A)** Schematic of approach used to determine how flattening of glucocorticoid oscillations affects adipocyte hypertrophy over time The perturbation from the normal glucocorticoid oscillation pattern is shown in orange. **(B)** Time course of average corticosterone levels in mice injected with corticosterone compared to mice injected with solvent control (phosphate buffer) 7 days after start of injections. n = 3-4, mean ± SEM. **(C)** Time course showing body weight gain of mice injected with corticosterone compared to mice injected with solvent control (phosphate buffer). n = 9 for sham injection; n = 9 for corticosterone injection, mean ± SEM. **(D)** Average daily food intake of each mouse per cage (n = 7). Up to 5 mice were housed in each cage. Food intake was monitored for 21 days, mean ± SD. Unpaired t test, **p < 0.01, ns = not significant. **(E-G)** sWAT, vWAT, and BAT fat pad weight measured at 3, 7, 14, and 21 days after start of daily injections, n = 3-8, mean ± SD. See also Table S1 and Figure S1.

Taken together, our results show that increasing the peak amplitude of corticosterone does not affect body weight gain or lipid accumulation in the white fat depots. Thus, as previously shown (Bahrami-Nejad et al., 2018), short spurts of glucocorticoids do not promote weight gain as long as the increase in glucocorticoids occurs during the time window that glucocorticoids are normally high and sufficiently long off-periods with low glucocorticoid levels are maintained. Furthermore, body weight gain and lipid accumulation in WAT and BAT were only observed in the cort-pellet mice with flattened glucocorticoids, despite the same total amount of glucocorticoid being introduced into both cort-pellet and cort-injected mice, arguing that it is the glucocorticoid flattening and not the amount of added glucocorticoids that is responsible for the adipocyte hypertrophy.

### Flattening of daily glucocorticoid oscillations does not result in fat deposition in the liver or increased fatty acids or triglycerides in the circulation

Chronic or misregulated glucocorticoid secretion has been shown to increase fat mass in rodents and humans, and the expansion of adipose tissue depots is accompanied by fat deposition in the liver and an increase of triglycerides and fatty acids in blood plasma (Bowles et al., 2015; Campbell et al., 2011; Chimin et al., 2014; Harris et al., 2013; Macfarlane et al., 2008). However, most *in vivo* studies administrated high pharmacological doses of glucocorticoids or administered glucocorticoids for several months which does not provide insight into how variations in normal physiological levels of glucocorticoids of limited duration – such as are seen in chronic stress, jet lag, night shift work - affect body metabolism. Since our experimental conditions use physiological levels of glucocorticoids and a limited three-week period of glucocorticoid flattening, we tested to see if there is still fat accumulation in liver. Markedly, histology analysis of the livers from cort-pellet mice showed no increase in lipid accumulation or fat pad weight (Figure 3A), or upregulation of lipid metabolism genes in the liver (Figure S2). Taken together, these results support that metabolic health is not yet strongly impacted in other tissues other that fat even after 3 weeks of glucocorticoid flattening.

**Figure 3.**
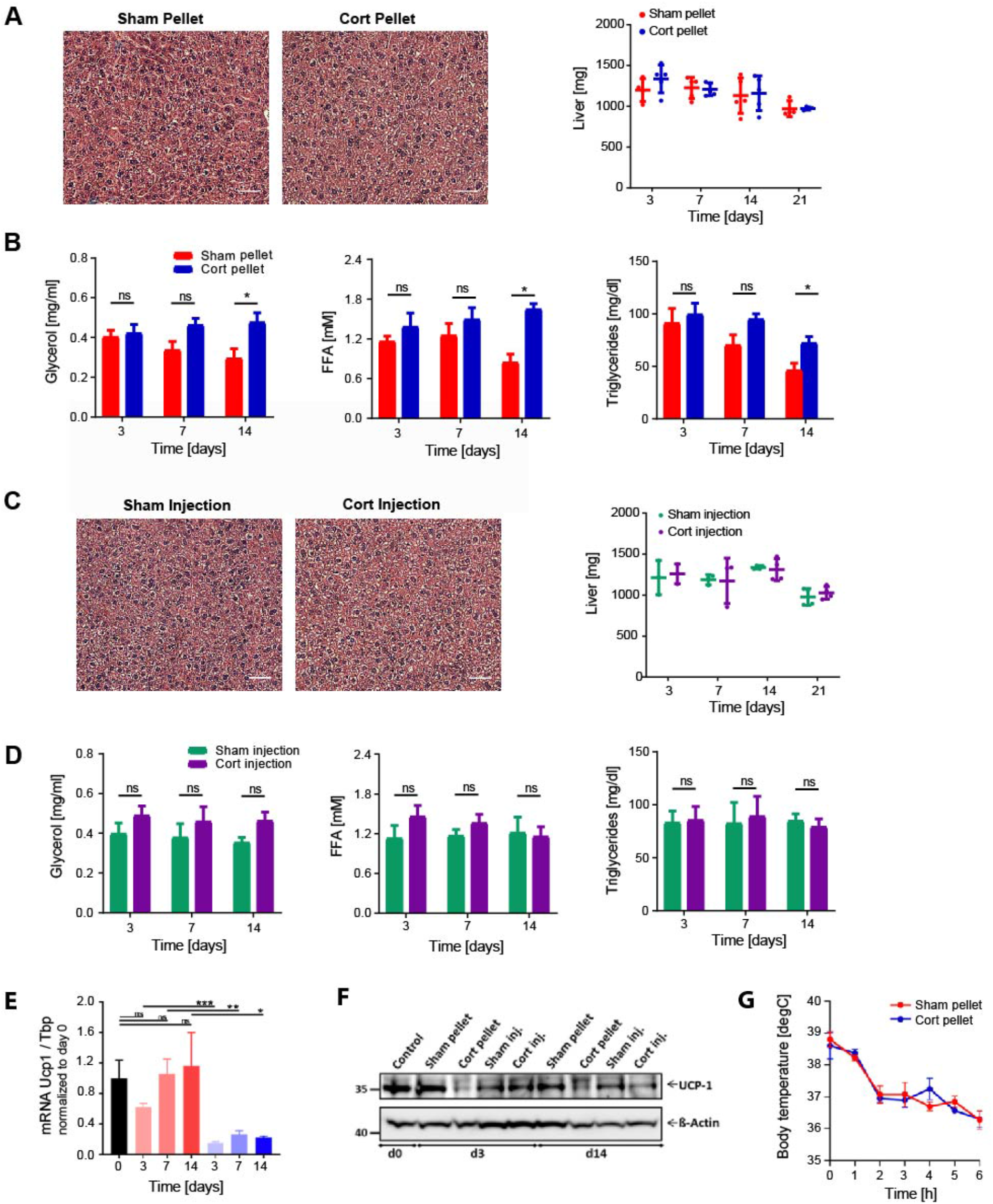
Flattening of glucocorticoid oscillations does not result in fat deposition in the liver, elevated circulating triglycerides in blood serum, or defects in thermogenesis. **(A)** H&E staining of livers 21 days after sham or corticosterone pellet implantation. One representative staining is shown (n = 4). Weight of livers at different time points after pellet implantation, n = 4-5, mean ± SD. See also Table S1. **(B)** Time courses of changes in FFA, glycerol, and triglycerides measured in blood serum over 14 days. n = 4-5 for pellet-implanted mice, mean ± SEM, unpaired t test, *p < 0.05, ns = not significant. **(C-D)** Same as in **(A-B)** but instead for daily Sham or corticosterone injected mice, n = 3-4. **(E)** Flattening of daily corticosterone oscillations in mice results in downregulation of genes involved in thermogenesis. mRNA expression for UCP-1 and PGC1a measured by qPCR in BAT of pellet implanted mice. Gene expression data was normalized to the expression of Tbp and is presented as mean ± SEM (n = 3-4). Unpaired t test, *p < 0.05, **p < 0.01, ***p < 0.001, ns = not significant. Control group are mice at start of experiment (day 0). **(F)** Western Blot analysis of UCP-1 in BAT lysates. **(G)** Acute cold exposure experiment of sham and cort pellet-implanted mice 21 days after pellet implantation. Mean +/- SEM; n = 8.

In another test of whether glucocorticoid flattening reduces overall metabolic health, we checked for increases in circulating triglycerides, or glycerol and free fatty acids (FFA), which are the building blocks of triglycerides. Given that triglycerides in adipose tissue are thought to be generated primarily from fatty acids taken up from the circulation rather than from *de novo* synthesized fatty acids (Santoro et al., 2021; Strawford et al., 2004), we expected to find that the adipocyte hypertrophy in WAT and BAT was caused by higher fatty acid levels in the circulation. Surprisingly, we did not detect significant differences in circulating glycerol, free fatty acids, or triglycerides between cort-pellet and sham-pellet mice after 3 and 7 days of flattening glucocorticoid oscillations (Figure 3B). Furthermore, what appeared as a small increase in these factors at day 14 was due primarily to reduced levels in controls. Thus, increased fatty acid levels in the plasma are not responsible for the rapid initial phase of adipocyte hypertrophy in observed in BAT and WAT during the first 2 weeks after glucocorticoid flattening. These results suggest that the initial increase in adipocyte hypertrophy is not yet associated with a loss of metabolic health. In control experiments, we confirmed that no significant lipid accumulation in the liver (Figure 3C) or changes in triglyceride, glycerol, and free fatty acid levels (Figure 3D) were detected when the same amount of corticosterone or control vehicle was injected during the glucocorticoid peak period.

### The expansion of WAT and BAT is not due to defective thermogenesis

Because we saw no lipid accumulation in the liver or significant lipid changes in the circulating blood plasma, we concluded that the adipocyte hypertrophy must be originating from an adipose tissue-intrinsic mechanism. It has been proposed that reduced thermogenesis due to defects in brown fat function can indirectly contribute to white fat mass enlargement (Strack et al., 1995; VanDenBeukel et al., 2014; Viengchareun et al., 2001). We thus focused next on asking whether a loss in brown fat thermogenesis could indirectly explain the rapid lipid accumulation in WAT. We used RT-PCR (Figure 3E) and western blot analysis (Figure 3F) to monitor UCP1 mRNA and protein expression in BAT of pellet-treated mice. Indeed, we did find reduced UCP1 in cort-pellet versus sham pellet mice, which is expected to contribute to reduced thermogenesis. However, when the mice were placed at 4 degC in an acute cold exposure experiment to compare their thermogenic capacities, we did not observe significant differences in how core body temperatures dropped (Figure 3h), supporting that the increased fat mass in the cort-pellet mice is not due to reduced thermogenesis despite reduced UCP1 expression in BAT (Luijten et al., 2019).

Taken together, the results so far suggest that adipocyte hypertrophy in response to glucocorticoid flattening is not caused by changes in the liver, plasma lipid levels or thermogenesis, suggesting that other metabolic changes in WAT and BAT are responsible. Since the primary role of glucocorticoids is to regulate gene transcription, we next performed a transcriptome time-course analysis of WAT and BAT tissues to understand whether transcription changes may cause the development of acute adipocytes hypertrophy in response to glucocorticoid flattening.

### Flattening of glucocorticoid oscillations results in strong and rapid upregulation of de novo fatty acid synthesis genes in WAT and BAT

We set out to compare differences in the adipocyte transcriptomes between mice that had oscillating versus flattened glucocorticoid levels. We focused on measuring gene expression in visceral white adipose tissue (vWAT) since its expansion has been recognized to be detrimental to metabolic health, while expansion of subcutaneous white adipose tissue (sWAT) may be protective (Ghaben and Scherer, 2019). We also analyzed brown adipose tissue (BAT) because of increasing evidence that BAT activity can significantly affect whole-body energy expenditure and fat tissue mass (Cannon and Nedergaard, 2004; Saito, 2013). We implanted mice subcutaneously with corticosterone or sham pellets and harvested vWAT and BAT between 1-5PM at four timepoints: 0, 3, 7, and 14 days after pellet implantation.

We carried out RNAseq analysis of vWAT and BAT in which we determined significant differential gene expression by an FDR corrected p-value (qval) less than 0.05. Using this cutoff criterion, we found several thousand transcripts were significantly changed in both vWAT and BAT when glucocorticoid oscillations were flattened (cort-pellet versus sham-pellet mice) (Figure S3 and Tables S4-S7). While the cort-pellet and sham-pellet samples were distinguishable from each other using PCA analysis (Figure 4A and Figure S4A), the samples obtained at different timepoints (3, 7 or 14 days) in the two treatment groups were largely indistinguishable from each other, supporting that flattening of glucocorticoid oscillations reprograms the vWAT and BAT transcriptomes within 3 days and that the changes are sustained for at least 2 weeks thereafter. That significant changes in several thousand genes of cort-pellet mice were already observed in vWAT at day 3 whereas expansion of vWAT was only observed at day 7 (Figure 1F), suggests that the glucocorticoid-mediated changes in the vWAT transcriptome help to drive the adipocyte hypertrophy in the vWAT, and not the other way around. Because the gene expression changes at 3, 7, or 14 days were largely indistinguishable from each other (Figure 4A), we combined the vWAT data from the three time points (3, 7, and 14 days) in analysis of gene expression changes in vWAT after glucocorticoid flattening. Similarly, we found for the BAT that gene expression changes at 3, 7, or 14 days in glucocorticoid flattened mice were largely indistinguishable from each other (Figure S4A), and thus, we also combined the BAT data from the three timepoints in our analysis of gene expression changes in BAT after glucocorticoid flattening.

**Figure 4.**
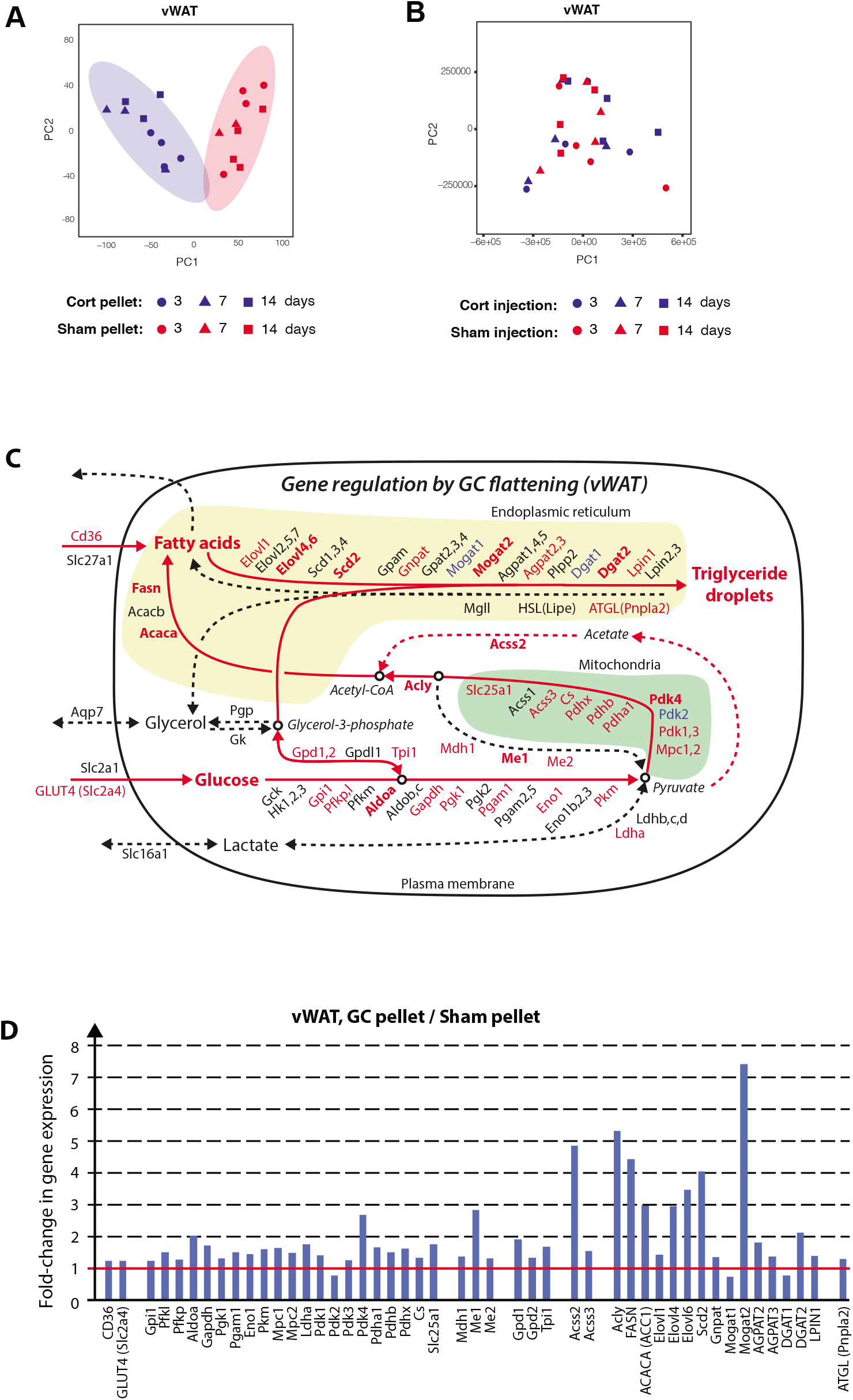
Flattening of glucocorticoid oscillations results in gene expression alterations in vWAT of genes involved in lipogenesis. **(A)** Principal component analysis of vWAT from corticosterone versus sham pellet-implanted mice obtained at 3, 7, and 14 days after pellet implantation. **(B)** Principal component analysis of vWAT from corticosterone and sham injected mice obtained at 3, 7, and 14 days after the start of injections. **(C)** Schematic of gene expression changes in triglyceride synthesis pathways in vWAT from corticosterone versus sham pellet-implanted mice. Upregulated, unchanged, and downregulated genes are marked in red, black, and blue, respectively. Bold type mark genes with a more than 2-fold change in gene expression. **(D)** Barplots showing fold-change of significantly changed vWAT genes in corticosterone versus sham pellet-implanted mice. See also Figures S4-S5 and Tables S2-S7.

In control experiments, we carried out RNAseq analysis of cort-injected and sham-injected mice. As mentioned before, these mice were injected at 5PM each day with the same amount of glucocorticoid released by the pellet per day. We sacrificed groups of mice between 1-5PM at four timepoints: 0, 3, 7 or 14 days after the start of daily injections and collected and analyzed vWAT and BAT by RNAseq. Remarkably, in comparing cort-injected versus sham-injected mice, PCA analysis showed no significant differences between the two conditions (Figure 4B and Figure S4B). Furthermore, no transcripts were changed in vWAT and only 9 transcripts were changed in BAT between cort-injected and sham-injected mice compared to the thousands of significant transcript changes observed in the vWAT and BAT between cort-pellet versus sham-pellet mice (Figure S3). Thus, even a large increase in the peak amplitude of corticosterone does not manifest as a significant change in gene expression in vWAT or BAT, consistent with our results in Figure 2 that showed no increase in fat mass in the cort-injected or sham-injected mice. Interestingly, since the time between injection of corticosterone at 5PM the day before until the mice were sacrificed between 1-5PM was between 20-24 hours, the fact that there are no significant changes in gene expression in the vWAT and BAT of cort-injected mice supports that the effect of glucocorticoids on gene expression is short-lived – less than 24 hours - in adipose tissue.

As an additional control, to validate that daily injection and sham pellets do not cause stress on their own that triggers gene expression changes, we compared sham-injected and sham pellet-implanted mice. No differently expressed genes were observed when comparing vWAT or BAT of these mice (Figure S3A-S3B), demonstrating that the procedure of pellet implantation and injections themselves do not alter gene expression and supporting that the injection and pellet-mediated delivery of glucocorticoid results can be directly compared.

Gene ontology (GO) analysis of the significantly regulated genes in vWAT of glucocorticoid flattened mice identified “fatty acid biosynthetic process”, “acetyl-CoA metabolic process”, “pyruvate metabolic process” and “glucose 6-phosphate metabolic process” as GO terms with high z-scores in vWAT and BAT (Tables S2 and S3). In addition, GO terms were enriched that were related to lipid uptake and utilization such as “lipid localization”, “fatty acid transport”, and “fatty acid metabolic process”. Similar metabolic GO terms were also identified in BAT. This analysis suggested that changes in adipocyte glucose and fatty acid metabolism may both contribute to the strong expansion of the BAT and WAT mass.

To better understand how flattening of glucocorticoid oscillations could be changing glucose and fatty acid metabolism in vWAT and BAT, we mapped our RNAseq data onto literature-curated maps that position metabolic genes into the main pathways that contribute to the synthesis and breakdown of triglycerides (Figure 4C and Figure S4). We colored each gene name by whether the transcript is significantly increased (red), unchanged (black) or decreased (blue). A gene was considered unchanged if the fold change was between 0.8 to 1.2 (less than 20% change). We used bold type for the gene name if the increase or reduction is greater than 2-fold.

Due to the known catabolic role of glucocorticoid, we had initially expected that flattening glucocorticoid oscillations would cause upregulation of the core lipolytic genes *Pnpla2* (*Atgl)*, *Lipe (Hsl)*, and *Mgll.* However, these genes were either weakly or not upregulated in BAT or vWAT. In contrast, we observed a striking upregulation of rate-limiting enzymes in the de novo fatty acid synthesis pathway such as fatty acid synthase (*Fasn*), Acetyl-CoA carboxylase (*Acaca and Acacb*), and *Acly*. We were surprised by this for two reasons. First, de novo fatty acid synthesis from glucose is not generally believed to be a main contributor to triglyceride synthesis in vWAT. Rather the main source of fatty acids to make triglycerides in adipocytes is thought to be fatty acids taken up from the circulation (Santoro et al., 2021; Strawford et al., 2004). Second, expression of lipogenesis genes such as Fasn and Acaca are known to be regulated by insulin signaling (Krycer et al., 2020; Song et al., 2018). This raised the question whether the transcriptome changes and adipocyte hypertrophy in mice with flattened glucocorticoid oscillations may at least in part be due to an increase in the level of insulin in the blood plasma. Previous studies, for example, showed a link from insulin signaling to glucose anabolism in adipocytes (Krycer et al., 2020).

Notably, there was a smaller upregulation of de novo fatty acid synthesis genes in BAT (Figure S4C-S4D). Instead, BAT has a stronger upregulation of *Cd36* expression compared to vWAT and an over 14-fold upregulation of *Gck* which is thought to be important in capturing glucose in cells (Matschinsky and Wilson, 2019), suggesting that BAT may use a different strategy to increase fat mass than vWAT when glucocorticoids are flattened and offering a possible explanation for the more rapidly observed adipocyte hypertrophy in BAT versus WAT (Figure 1G),

### Glucocorticoid flattening results in rapidly induced hyperinsulinemia

At the outset of our study, we did not consider that insulin might be elevated in the cort-pellet since insulin is short-lived in the circulation and high or misregulated glucocorticoids levels are commonly thought to inhibit insulin release from pancreatic beta cells (Delaunay et al., 1997). However, we realized that hyperinsulinemia is common in human patients with flattened circulating glucocorticoid levels such in subcritical hypercortisolism (Elhassan et al., 2019; Lopez et al., 2016; Rossi et al., 2000). Furthermore, other studies have shown that hyperinsulinemia, even when glucose was low, could originate from defects in insulin clearance (Bojsen-Møller et al., 2018; Protzek et al., 2016)and also that glucocorticoid-stimulated pancreatic beta cells can in some cases still secrete insulin (Fine et al., 2018). Taken together, it was plausible that high insulin could be responsible for the gene expression profiles we had measured in mice with flattened glucocorticoid oscillations. We thus carried out a timecourse analysis in which we monitored insulin levels in blood plasma over 21 days in the cort-pellet and sham-pellet mice.

Strikingly, we found that in mice with flattened corticosterone oscillations insulin levels increased within 3 days to over 7 ng/ml compared to less than 1 ng/ml in sham pellet-implanted mice (Figure 5A). This unexpectedly initial high insulin level then slowly decreased during the 21-days of glucocorticoid flattening to about 3 ng/ml, which was still several fold higher than the insulin levels in sham pellet-implanted mice. High insulin levels in mice and humans are typically correlated with high glucose levels, such as occur after a meal, that stimulate pancreatic beta cells to secrete more insulin. We were thus surprised to find that glucose levels are low in glucocorticoid-flattened mice, even slightly lower than in control mice (Figure 5B). Since glucose levels are not high, this suggests that it is not low glucose that signals to pancreatic beta cells to secrete more insulin. As alternative mechanisms, glucocorticoids have been shown to reprogram pancreatic beta cells to cause continued insulin secretion by increasing cAMP levels (Fine et al., 2018), as well as to cause decreased hepatic clearance of insulin (Protzek et al., 2016).

**Figure 5.**
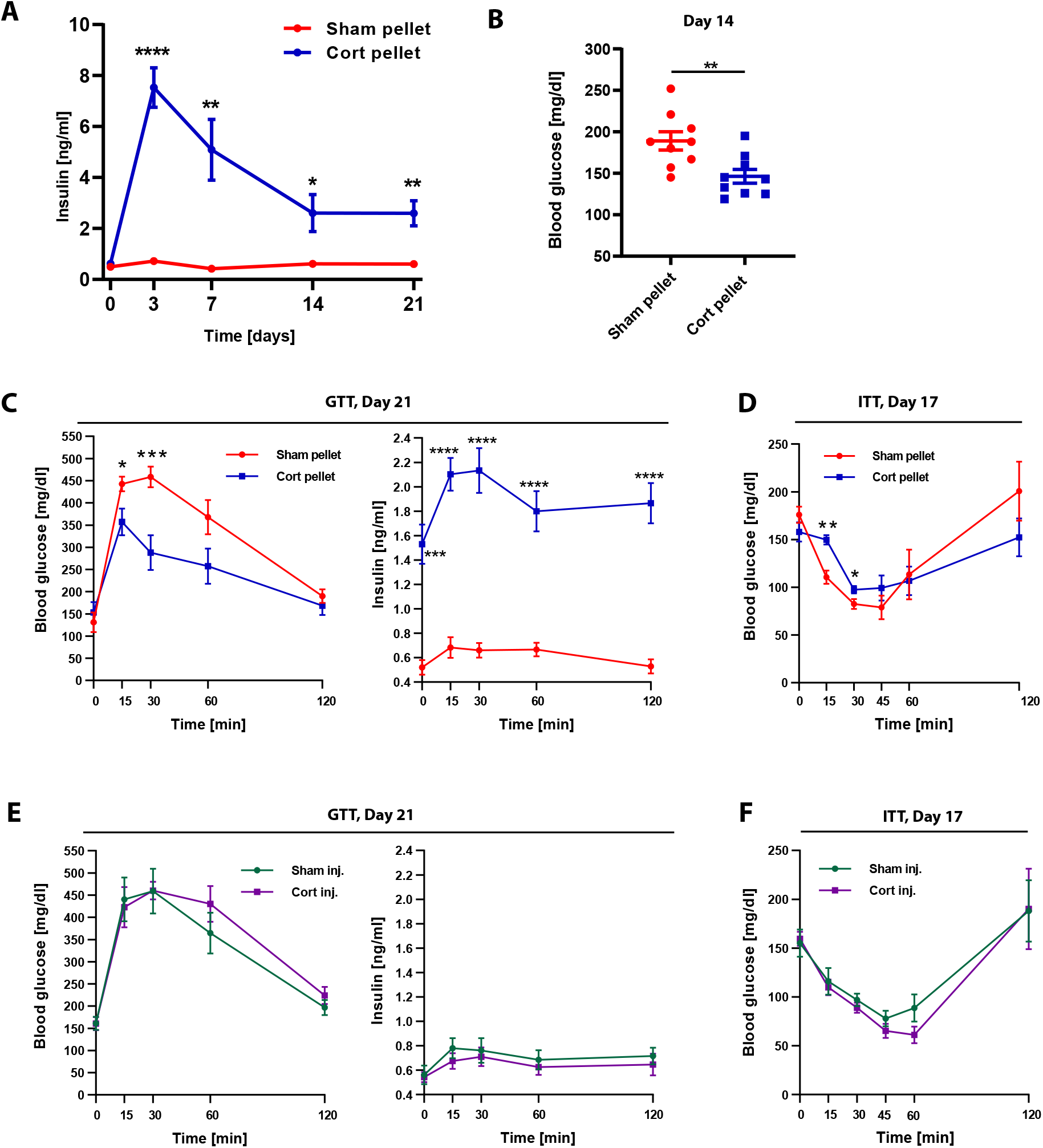
Flattening of glucocorticoid oscillations causes hyperinsulinemia within 3 days. **(A)** Time course of insulin levels in blood plasma in sham and corticosterone pellet-implanted mice (not fasted); mean +/- SEM; n = 6; unpaired t test, *p < 0.05, **p < 0.01, ****p < 0.0001. **(B)** Blood glucose levels in sham and corticosterone pellet-implanted mice (not fasted) 14 days after pellet implantation; mean +/- SEM; n = 9; unpaired t test, **p < 0.01. **(C)** Glucose tolerance test (GTT) of sham and corticosterone pellet-implanted mice after 21 days of treatment; mean ± SEM; n = 5; unpaired t test, *p < 0.05, ***p < 0.001, ****p < 0.0001. **(D)** Insulin tolerance test (ITT) of sham and corticosterone pellet-implanted mice after 17 days of treatment; mean ± SEM; n = 5 for sham pellet; n = 6 for corticosterone pellet; unpaired t test, *p < 0.05, **p < 0.01. **(E)** Glucose tolerance test (GTT) of sham and corticosterone injected mice after 21 days of treatment; mean ± SEM; n = 6. **(F)** Insulin tolerance test (ITT) of sham and corticosterone injected mice after 17 days of treatment; mean ± SEM; n = 5 for sham injection; n = 6 for corticosterone injection.

Given the persistent high levels of insulin, we also considered that high insulin for continuous periods of days and weeks may result in a loss of insulin sensitivity or ability to properly clear glucose from the bloodstream. Thus, we carried out insulin and glucose tolerance tests (ITT and GTT) at timepoints when insulin had been high for 17-21 days. Interestingly, as shown in the GTT test results (Figure 5C), an intraperitoneally injected bolus of glucose was taken up even more rapidly in glucocorticoid pellet-implanted mice compared to sham pellet control mice. The faster drop in glucose seemed initially surprising but likely results from the higher level of insulin in the glucocorticoid pellet implanted mice.

The fast clearance of a bolus of glucose in the GTT tests suggested that the glucocorticoid flattened mice are still insulin sensitive. To test whether the mice are also still insulin sensitive despite glucocorticoid-flattening and hyperinsulinemia, we carried out ITTs. We injected boluses of insulin into the mice and measured whether glucose levels would drop within 30 minutes (Figure 5D). Indeed, glucose levels in glucocorticoid flattened mice dropped similarly as in the control mice, albeit slightly slower, supporting that corticosterone pellet-implanted mice are still insulin sensitive but have a weak form of insulin-resistance. The hyperinsulinemia when glucocorticoid oscillations are flattened, together with the maintenance of insulin sensitivity, can explain the observed slightly lower-than-normal glucose levels since insulin promotes glucose uptake into fat and muscle ^43^. Interestingly, the lower blood glucose levels (Figure 5B), and thus likely lower metabolism in the glucocorticoid-flattened mice, provides an explanation for why these mice are able to gain weight, even though we found no change in food intake during 3 weeks of glucocorticoid flattening (Figure 1D).

In control experiments, mice injected with the same dose of glucocorticoid as released by the pellet per day did not reveal any differences in glucose tolerance compared to sham injected mice and did not display higher insulin levels in the blood plasma (Figures 5E-5F). Together, these results show that flattening of glucocorticoids causes hyperinsulinemia that occurs within 3 days, before adipocyte hypertrophy is observed in WAT.

### Metabolic changes caused by 3 weeks of flattening glucocorticoids are largely reversible

One of the goals of this study was to mimic a temporary three week-long stress condition and accordingly chose a pellet that releases corticosterone over 3 weeks. During the three weeks of glucocorticoid flattening, we observed hyperinsulinemia and significant adipocyte hypertrophy which are typically considered to be harmful for metabolic health. It was therefore surprising that mice were able to maintain low glucose levels and insulin sensitivity, which are markers of a healthy metabolic state, for the three weeks in which glucocorticoid were flattened. Furthermore, the levels of fatty acids and triglyceride in the circulation were only minimally increased and no lipid accumulation was observed in the liver (Figure 3), further suggesting that mice in which glucocorticoids are flattened even for up to 3 weeks are still metabolically healthy, despite significant weight gain, adipose tissue hypertrophy, and hyperinsulinemia.

Strikingly, the hyperinsulinemia in glucocorticoid flattened mice was fully reversible. Insulin levels returned to the levels of the sham-pellet mice at 42 days, which is 3 weeks after the time when the implanted pellet stops releasing corticosterone (Figure 6A). These results support a direct role for flattening of glucocorticoid oscillations in driving the hyperinsulinemia. Furthermore, about half of the weight gain in WAT and all the weight gain in BAT were lost at 42 days (Figure 6B). There was variability between mice, and many restored their fat mass to that of control mice at that time. Together, these results suggest that the greater than two-fold increase in adipocyte hypertrophy is mostly reversible in WAT and fully reversible in BAT over a similar time period as it took to increase the fat mass. However, it should be noted that while the healthy low trough glucocorticoid levels were restored by 3 weeks after the pellet stopped releasing glucocorticoid (Figure S5), the peak levels during waking remained reduced in most of the mice, suggesting that the 3-week glucocorticoid flattening has a longer-term impact on the diurnal control of glucocorticoid secretion from the adrenal cortex.

**Figure 6.**
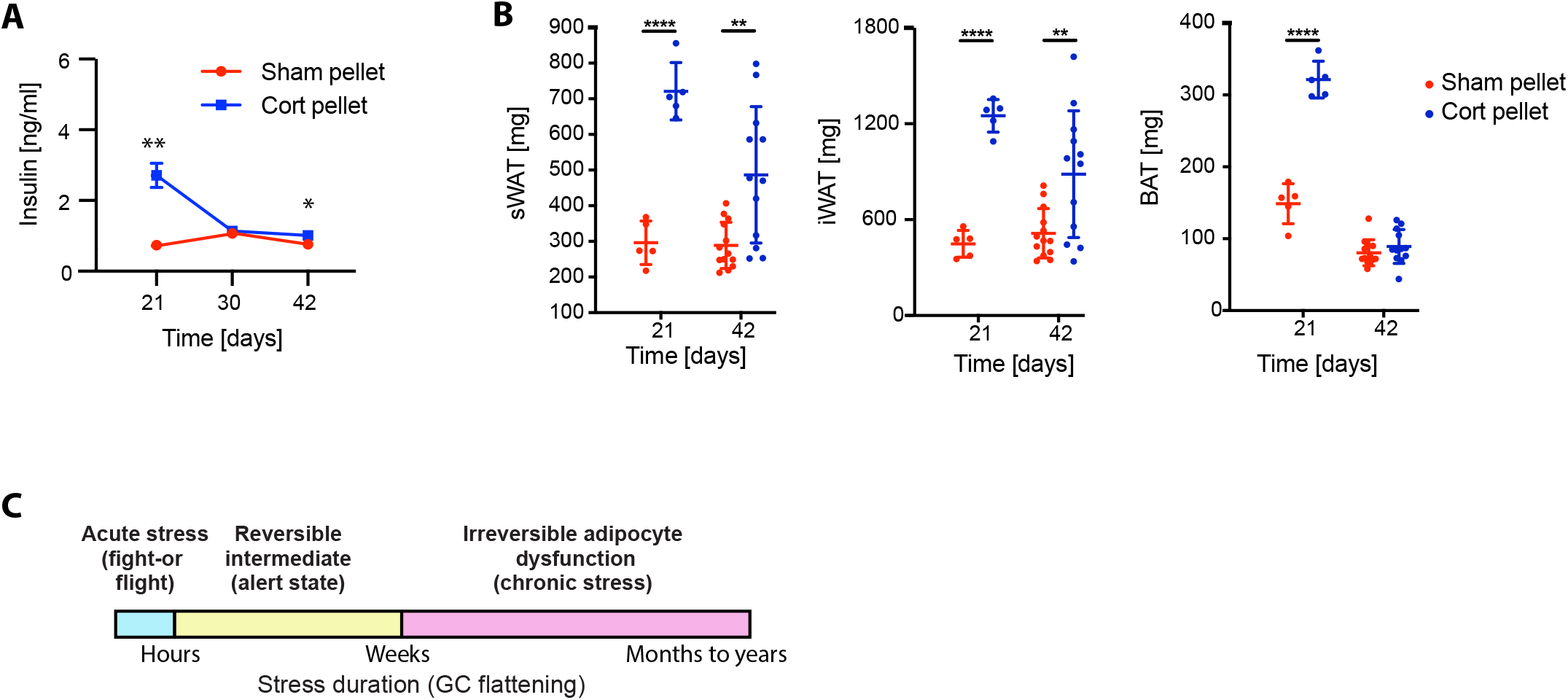
The hyperinsulinemia and much of the adipocyte hypertropy are reversible when the glucocorticoid flattening stimulus is removed. **(A)** Time course of insulin levels in blood plasma in sham and corticosterone pellet-implanted mice (not fasted); mean +/- SEM; n = 6; unpaired t test, *p < 0.05, **p < 0.01. **(B)** Fat pad weight was measured at 21 and 42 days after pellet implantation, n = 5-9, mean ± SD. Unpaired t test, **p < 0.01, ****p < 0.0001. Data from 21 days is the same as in Figures 1E-1F, second panel. **(C)** Schematic representation of the commonly discussed acute and chronic stress states, as well as a reversible intermediate state arising from three weeks of glucocorticoid flattening. Mice can mostly recover from this intermediate alert state without long-term adipocyte dysfunction.

We conclude that glucocorticoid flattening rapidly induces white and brown adipocyte hypertrophy, causing WAT and BAT to more than double in only three weeks, but the increase in fat mass remains mostly reversible and the hyperinsulinemia remains fully reversible. These results support that there is a healthy “alert” state of intermediate duration that mice can assume for weeks in response to temporary stress and continually activated glucocorticoid signaling. This alert state has a protective role by keeping the levels of fatty acid and glucose in the circulation in check due to high circulating insulin and high glucose uptake rates. The existence of this alert state is intriguing since most previous work has described stress as being either acute, fight-or-flight response glucocorticoid increases or chronic long-term stress that would cause irreversible adipocyte dysfunction and associated metabolic diseases (McEwen, 2000; Sapolsky, 2004; Sapolsky et al., 1986). Before this study, our assumption had been that adipocyte hypertrophy caused by a chronic three-week long glucocorticoid flattening would be irreversible. However, our data shows that this alert state is reversible, and mice can recover from the hyperinsulinemia and adipocyte hypertrophy, as long as the stress and glucocorticoid flattening does not exceed a few weeks. Taken together, these results suggest that there should be a modification to the commonly discussed distinction between an acute, fight-or-flight type glucocorticoid response on the order of hours, and a chronic permanent glucocorticoid dysregulation (McEwen, 2000; Sapolsky, 2004; Sapolsky et al., 1987). Our study suggests that one should add a reversible alert state of intermediate duration, approximately of 3 weeks. Based on previous studies that used longer glucocorticoid flattening (Dunford and Riddell, 2016; Lee et al., 2018), this intermediate state would then transition to a chronic irreversible state if the stress lasts much longer or glucocorticoid signals are elevated (Figure 6C, yellow box).

### CD36ko mice are partially protected against weight gain induced by flattening of glucocorticoid oscillations

The hyperinsulinemia (Figure 5) and large metabolic transcriptome changes in WAT and BAT (Figure 4) are already fully induced within three days of flattening of glucocorticoid oscillations, which is before expansion of WAT depots is observed. Since insulin is known to mediate many of the transcriptome changes observed after 3 days, a likely order of events is that mice with flattened glucocorticoid rapidly develop hyperinsulinemia which then causes insulin-driven upregulation of fatty acid synthesis genes such as *Fasn, Acaca,* and *Gck* in fat cells. Upregulation of these genes would increase conversion of glucose into fatty acids and increase triglyceride synthesis, explaining the adipocyte hypertrophy observed a few days later.

We nevertheless considered that the adipocyte hypertrophy could also be partially due to increased fatty acid uptake, especially since Cd36, a high-affinity receptor for long-chain fatty acids that facilitates fatty acid uptake into muscle and adipose tissues of rodents and humans(Pepino et al., 2014), has been shown to be glucocorticoid-regulated (Yu et al., 2010). Our RNAseq analysis had shown that CD36 gene expression levels are upregulated in both vWAT and BAT of cort-pellet mice (Figures 4C, 4D, S4C, and S4D). RT-PCR analysis confirmed that Cd36 was indeed upregulated (Figure 7A), We thus went forward with investigating CD36’s role in adipocyte hypertrophy when glucocorticoid oscillations are flattened.

**Figure 7.**
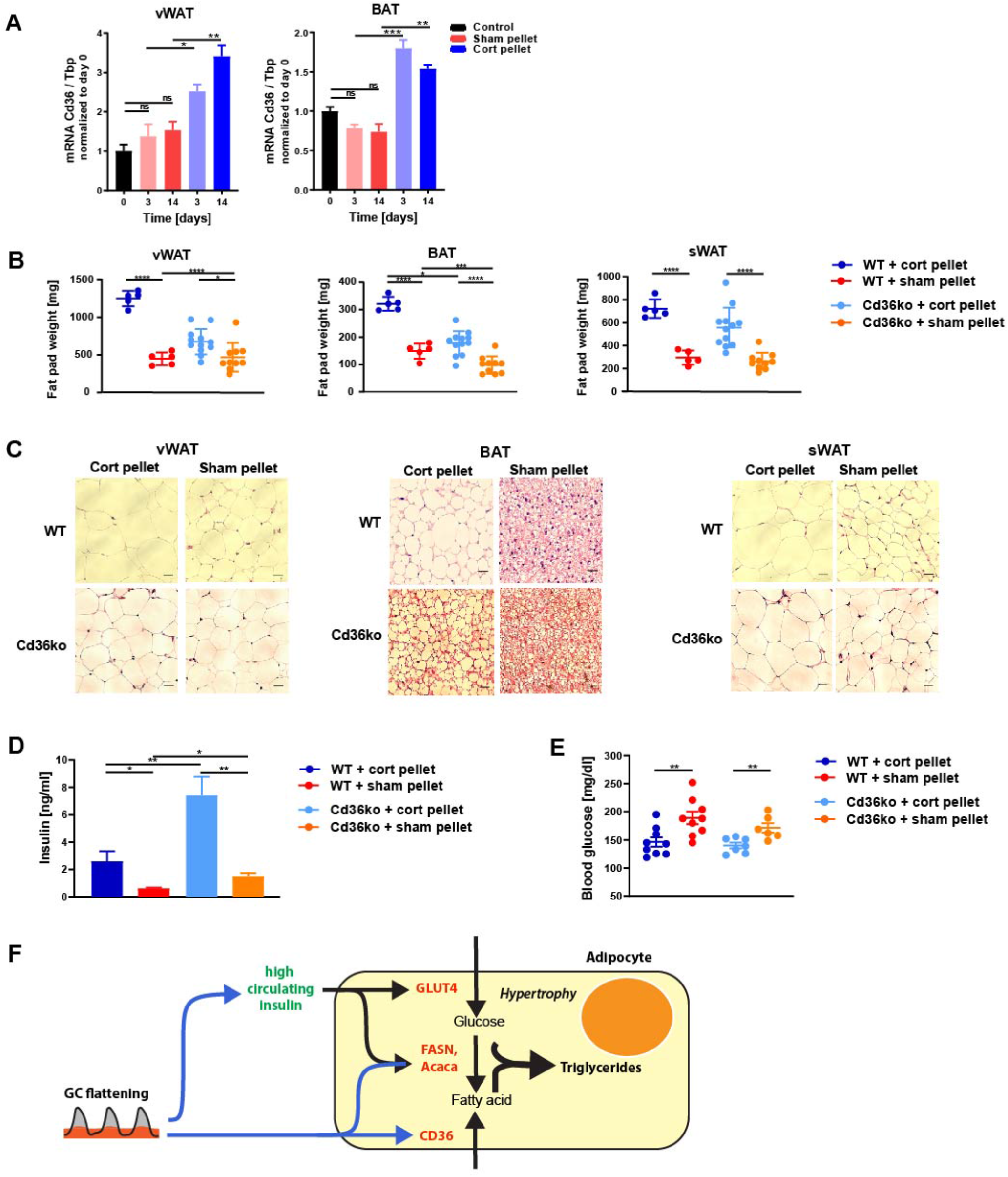
Cd36ko partially protects against adipocyte hypertrophy resultng from flattening of glucocorticoid oscillations. **(A)** qPCR of vWAT and BAT 3 and 14 days after cort or sham pellet implantation. Data is normalized to the expression of Tbp and is presented as mean ± SEM (n = 3-4). Unpaired t test, *p < 0.05, **p < 0.01, ***p < 0.001, ns = not significant. Control group are mice at start of experiment (day 0). **(B)** Weight of vWAT, BAT, and sWAT fat depots 21 days after pellet implantation, n = 5-12, mean ± SD. Unpaired t test, *p < 0.05, ****p < 0.0001. See also Table S1. **(C)** H&E staining of vWAT, BAT, and sWAT 21 days after pellet implantation. One representative staining is shown (n = 3-5). Scale bars 50µm. See also Figure S6. **(D)** Insulin levels in blood plasma in corticosterone and sham pellet-implanted Cd36ko and wild-type mice (not fasted) 14 days after pellet implantation; mean +/- SEM; n = 5-7; unpaired t test, *p < 0.05, **p < 0.01. **(E)** Blood glucose levels in corticosterone and sham pellet-implanted Cd36ko and wild-type mice (not fasted) 14 days after pellet implantation; mean +/- SEM; n = 6-9; unpaired t test, **p < 0.01. **(F)** Scheme of how an initially protective adipocyte hypertrophy mechanism can be activated in response to flattening of circadian glucocorticoid oscillations. Glucocorticoid flattening results in a unique metabolic signature with high insulin, slightly lower-than-normal glucose, and normal FFA levels in the circulation.

We implanted Cd36ko and wildtype mice with corticosterone or sham pellets for 21 days and then dissected and weighed their sWAT, vWAT and BAT tissues. As had been observed previously in Figures 1E-1G, flattening glucocorticoid oscillations in wildtype mice increased vWAT, sWAT and BAT weight more than two-fold over control mice (dark blue compared to red; Figure 7B). Remarkably, knocking out Cd36 resulted in a much smaller increase in weight for all three fat tissue types (light blue compared to orange; Figure 7B). Histological analysis of the three fat tissue types showed that adipocytes of cort-pellet Cd36ko mice displayed lower increases in cell volume than cort-pellet wildtype mice (Figure 7C; quantification is shown in Figure S6), further supporting that at least part of the adipose hypertrophy in glucocorticoid flattened mice is caused by Cd36-mediated fatty acid uptake.

Although loss of Cd36 significantly reduces the adipocyte hypertrophy in WAT and BAT upon flattening corticosterone oscillations, the cort-pellet Cd36ko mice still have hyperinsulinemia (Figure 7D) and lower blood glucose levels (Figure 7E), similar to the cort-pellet wildtype mice. That the rescue of adipocyte hypertrophy in Cd36ko mice is only partial is consistent with the interpretation that the remaining hypertrophy is the result of hyperinsulinemia. Together, these results suggest that adipocyte hypertrophy in response to glucocorticoid flattening is caused by a combination of Cd36-mediated uptake of fatty acids and hyperinsulinemia-mediated increases in glucose uptake and de novo fatty acid synthesis from glucose.

## DISCUSSION

Flattening of daily glucocorticoid oscillations in rodents and humans due to Cushing’s disease and chronic stress has been shown to cause obesity and metabolic disease (Bahrami-Nejad et al., 2018; Campbell et al., 2011; Chimin et al., 2014; Dallman et al., 2000; Joseph and Golden, 2017; Karatsoreos et al., 2010; Rossi et al., 2000), with adipocyte hypertrophy believed to be a main contributing factor. Here we sought to understand the mechanisms underlying the adipocyte hypertrophy by varying the timing of how glucocorticoids were applied to mice and measuring relevant parameters in blood and tissues at multiple timepoints. Strikingly, within 3 days, circulating insulin levels became persistently high and thousands of transcripts in WAT and BAT changed. These effects preceded the adipocyte hypertrophy and expansion of WAT observed at 7 days in the white fat (WAT).

Since adipocyte hypertrophy is typically associated with increased fatty acids and triglycerides in the circulation and increased fat deposition in the liver (Campbell et al., 2011; Harris et al., 2013), we were surprised to find no increase in circulating fatty acid levels and no measurable fat deposition in the liver. Furthermore, because hyperinsulinemia is typically closely associated with high glucose levels, we were also surprised to find that circulating glucose levels were even slightly lower in mice with flattened glucocorticoids and hyperinsulinemia, compared to control animals. An explanation for the low glucose levels is that even after 3 weeks of glucocorticoid flattening and significant hyperinsulinemia and adipocyte hypertrophy, the mice are still able to uptake glucose into peripheral tissues at even a higher rate compared to control mice, as demonstrated by a glucose tolerance test. These results suggests that the significant adipocyte hypertrophy observed in the glucocorticoid flattened mice is initially not causing a loss of metabolic health.

It has generally been assumed that most of the lipid stored in adipocytes is synthesized from fatty acids taken up from the circulation, rather than being synthesized from glucose (Digirolamo et al., 1992; Santoro et al., 2021). However, our analysis of CD36 knockout mouse showed that increased fatty acid uptake was only partially responsible for the adipocyte hypertrophy. For conditions in which insulin is high and glucose is low, as we observe in glucocorticoid flattened mice, glucose uptake is accelerated, and the significant upregulation of de novo fatty acid synthesis genes observed in vWAT and BAT in our RNAseq analysis support that the percentage of glucose converted to triglycerides is high. Furthermore, the lack of increase in lipolytic genes in vWAT and BAT and low circulating free fatty acids support that the newly synthesized triglycerides are retained in the adipocytes. Taken together, our results argue that increased insulin-mediated glucose uptake, de novo synthesis of fatty acids from glucose, and suppression of lipolysis in adipocytes drive the adipocyte hypertrophy in the glucocorticoid-flattened mice. The hyperinsulinemia we observed when glucocorticoids are flattened is consistent with observations in human patient studies of subcritical hypercortolism (Papanastasiou et al., 2017; Rossi et al., 2000). Further studies are needed to determine the cause of the hyperinsulinemia in the absence of increased glucose when glucocorticoids are flattened and whether it be due to increased insulin secretion from pancreatic beta cells or reduced insulin clearance in the liver requires further study (Bojsen-Møller et al., 2018; Protzek et al., 2016).

The implanted pellet was designed to release corticosterone at a constant rate for 21 days. Interestingly, analysis of mice at 42 days, three weeks after the glucocorticoid flattening stimulus ceased, showed that the hyperinsulinemia was completely reversed, glucocorticoid levels during the trough period had returned to basal levels, and about half of the gain in WAT mass and all of the gain in BAT mass were reversed as well. We further found normal levels of circulating FFAs and no lipid accumulation in the liver after 2-3 weeks of glucocorticoid flattening, supporting that there is no significant loss of metabolic health during this period, despite a more than doubling of fat mass and lipid accumulation in WAT and BAT. Nevertheless, there were signs for longer term effects on the overall metabolism in mice after 3 weeks of glucocorticoid flattening. Even after the CORT pellet had ceased to release corticosterone for 3 weeks, white fat mass in both vWAT and sWAT were still 50% higher and the normally high peak levels of corticosterone during waking were not yet fully restored. There was much variability between mice with some returning to normal fat mass levels but others keeping a higher mass. Nevertheless, taken together, despite the more than doubling of fat mass, our studies suggest that cells remain metabolically healthy up to a 3-week period of glucocorticoid flattening before flattening of glucocorticoids ends up causing metabolic disease and diabetes..

A plausible overall interpretation of our results is that glucocorticoid flattening leads to a metabolic “alert” state that is distinct from the initial rapid fight-or-flight response and distinct from the chronic stress state that is associated with metabolic diseases. This alert state is marked by persistently high insulin and increased Cd36-mediated fatty acid uptake into adipocytes that results in normal glucose and fatty acid levels in the circulation (Figure 7F). For periods of up to three weeks, a gradual increase in adipose hypertrophy helps maintain the alert state by accelerated uptake of glucose and fatty acid. The cost of maintaining this alert state for three weeks is a greater than two-fold increase in white and brown adipose mass due to continued uptake of fatty acids and glucose into WAT and BAT that ultimately results in metabolic disease.

In summary, our study shows the cellular and physiological consequences of temporary flattening of glucocorticoids that is frequently observed during chronic stress. We show that loss of the normal diurnal glucocorticoid oscillation pattern can result in acute hyperinsulinemia and acute hypertrophy in which white and brown fat mass more than doubles within 3 weeks. The rapid increase in white and brown fat mass is mediated by a combination of insulin-mediated upregulation of glucose conversion to fatty acid and Cd36-mediated fatty acid uptake. Remarkably, for up to about 3 weeks, fatty acids and glucose levels in the circulation remain near normal levels and much of the metabolic changes associated with poor metabolic health remain reversible. Furthermore, much of the gain in white fat mass and all of the gain in brown fat mass can be reversed when glucocorticoid flattening stops. Continued fatty acid and glucose uptake into fat tissue can therefore minimize the damage that could be caused by hyperglycemia and high lipid levels in the circulation resulting from glucocorticoid-mediated catabolism. Thus, to maintain the alert status for a potential fight-or-flight response, the hyperinsulinemia and adipocyte hypertrophy induced by glucocorticoid flattening are unexpectedly protective for metabolic health for a period of a few weeks.

## AUTHOR CONTRIBUTIONS

S.T. and M.N.T. conceived the project. S.T. performed and analyzed all experiments with assistance of A.R., E.B.M., and W.Y. K.M.K. analyzed transcriptomic data. F.B.K. designed experiments and contributed to the discussion. S.T. and M.N.T wrote the paper with input from all authors.

## ACKNOWLEDGEMENTS

This work was supported by National Institutes of Health RO1-DK101743, RO1-DK106241, P50-GM107615, and Stanford BioX Seed Grant funding (to M.N.T.), and the U.S. Department of Veterans Affairs I01BX000398 (FBK). Furthermore, we acknowledge the funding from the Deutsche Forschungsgemeinschaft (DFG) TH 2156/1-1 (to S.T.), the Stanford Bio-X Program and Novo Nordisk Foundation NNF16OC0018642 (to A.R.), and American Heart Association Postdoctoral Fellowship 18POST34030448 (to E.B.M.). We thank Tim McGraw, Fred Maxfield, Laura Alonso, Tobias Meyer (Weill Cornell Medicine) and the members of the Teruel Lab for discussions and critical reading of the manuscript. We also thank the Stanford Animal Histology Services (AHS) for paraffin embedding of fat tissue and performing H&E staining.

## Declaration of interest

The authors declare no competing interests.

## SUPPLEMENTARY FIGURES

**Figure S1. Images of tissue and adipocyte sections in WAT and BAT induced by daily corticosterone injection versus sham injection.**

**(A-C)** Lack of a visible change in size of WAT and BAT tissue and size of adipocytes in Cort injected versus Sham injected mice (n=3 mice per treatment, 25 images per mouse (20x)).

**Figure S2. Flattening of corticosterone oscillations does not result in significant gene expression alterations of genes involved in lipid metabolism in liver.**

**(A-J)** Gene expression of different genes measured by qPCR in liver after flattening corticosterone levels for 14 days. Gene expression data was normalized to the expression of Tbp and is presented as mean ± SEM (n = 3-4). Unpaired t test, *p < 0.05, **p < 0.01, ns = not significant. Control group are mice at beginning of experiment (day 0).

**Figure S3. Comparison of BAT and vWAT RNAseq datasets.**

**(A-B)** Number of significantly changed transcripts in BAT and vWAT in different comparisons of treatment methods.

**(C)** Number of overlapping transcripts in vWAT and BAT. See also Tables S2-S7.

**Figure S4. Analysis of BAT RNAseq dataset.**

(A) Principal component analysis of BAT from corticosterone versus sham pellet-implanted mice obtained at 3, 7, and 14 days after pellet implantation.

(**B**) Principal component analysis of BAT from corticosterone and sham injected mice obtained at 3, 7, and 14 days after the start of injections.

(**C**) Schematic of gene expression changes in triglyceride synthesis pathways in BAT from corticosterone versus sham pellet-implanted mice. Upregulated, unchanged, and downregulated genes are marked in red, black, and blue, respectively. Bold type mark genes with a more than 2-fold change in gene expression. See also Tables S2-S7.

(**D**) Barplots showing fold-change of significantly changed BAT genes in corticosterone versus sham pellet-implanted mice. See also Tables S2-S7.

**Figure S5. Corticosterone measurements in mice implanted with corticosterone or sham pellets at Day 42 after pellet implantation.**

**(A)** Blood serum corticosterone levels at Day 42 for 8 corticosterone pellet-implanted mice.

**(B)** Blood serum corticosterone levels at Day 42 for 8 sham pellet-implanted mice.

**Figure S6. Quantification of lipid droplet size in WT and CD36KO mice implanted with corticosterone or sham pellets for 21 days.**

**(A-C)** Frequency distributions of mean lipid droplet area in sWAT, vWAT, and BAT as analyzed using Adiposoft (mean + SEM; n=3 mice per treatment, 25 images per mouse (20x)).

## SUPPLEMENTARY TABLES

**Table S1. Summary of mouse experiments.** All data was acquired after a treatment period of 21 days, except data for blood glucose (not fasted), FFA, glycerol, and triglyceride levels, which were determined 14 days after beginning of treatment. Values are presented as mean +/- SEM; n.d. = no data.

**Table S2. Gene otology (GO term) analysis of differentially expressed genes in vWAT of corticosterone versus sham pellet-implanted mice.**

**Table S3. Gene otology (GO term) analysis of differentially expressed genes in BAT of corticosterone versus sham pellet-implanted mice.**

**Table S4. Transcript levels of differentially expressed genes in vWAT of corticosterone versus sham pellet-implanted mice.**

**Table S5. Gene expression levels of differentially expressed genes in vWAT of corticosterone versus sham pellet-implanted mice.**

**Table S6. Transcript levels of differentially expressed genes in BAT of corticosterone versus sham pellet-implanted mice.**

**Table S7. Gene expression levels of differentially expressed genes in BAT of corticosterone versus sham pellet-implanted mice.**

## METHODS

### Mice

Mice were housed in small groups of five or fewer on a 12h light/dark cycle (lights on at 7:00 hours) with food and water access ad libitum in the animal facility at Stanford University. All mice were maintained on a normal chow diet and only male mice were used for experiments. All animal care and experimentation was conducted in accordance with current NIH and Stanford University Institutional Animal Care and Use Committee guidelines. Seven-week-old C57BL/6J male mice were purchased from Jackson Laboratory (cat. 000664) and housed in the animal facility for 7 days prior the start of experiments. Cd36ko mice were published before(Febbraio et al., 1999). Cd36ko breeding pairs were purchased from Jackson Laboratory (cat. 019006). For the experiments, 8-week old male mice were used. All mice were dissected between 13:00 and 18:00 alternating corticosterone-treated and control mice. A summary of all parameters measured in mice upon different treatments can be found in Extended Data, Table 1.

During the acute cold exposure experiments, mice were placed in individual cages with minimal bedding in order to prevent nesting or huddling behavior which would interfere with adaptive thermogenesis. The cages were placed in a 4 °C deli fridge for the 6-hour timecourse experiment. Core body temperatures of the mice were assessed with a rectal probe thermometer (Type J/K/T Thermocouple meter, Digisense, Kentscientific, USA; with rectal probe for mice RET-3, Physitemp Instruments, USA) before and at one-hour intervals during the 6-hour cold exposure. Timepoint 0 was taken 1 minute before the mice went into the cold.

### Corticosterone administration in mice

To flatten diurnal glucocorticoid oscillations, mice were implanted subcutaneously with corticosterone releasing pellets (5mg, 21-day release; Innovative Research of America, Sarasota, FL, USA, Catalog number G-111). Sham pellets (Catalog number C-111) were implanted as control. For pellet implantation mice were anesthetized via inhalation of isoflurane, when anesthetized an incision equal in diameter to that of the pellet was made on the lateral side of the neck and the pellet inserted using a trochar. Mice weighed an average of 24.1 ± 1.2 g, which results in a daily dose of 9 mg/kg/day.

For injections, corticosterone complexed with 2-hydroxypropyl-β-cyclodextrin (C174, Sigma) was dissolved in phosphate buffer (PBS) and injected subcutaneously once daily at 17:00 for up to 21 days with the same corticosterone dose (9 mg/kg/day) as released by the corticosterone pellets per day. PBS was injected as control.

### Measurement of corticosterone in blood serum

Blood was taken at multiple time points over a 15 h time period. At the first time point, blood was taken by nicking the tail vain. Blood samples collected on following time points were taken by removal of the crust formed after first blood withdrawal. The blood was allowed to clot by leaving it undisturbed at room temperature for 45 minutes. The clot was removed by centrifuging at 2000 x g for 15 minutes. The corticosterone concentration in blood serum was determined using the Enzyme Immunoassay kit (K014-H1, Arbor Assays, Michigan, USA) following the manufacturer’s instructions.

### Glucose Tolerance Test

Before the procedure mice were fasted for 16h in clean cages with no food or feces in the bedding. At 9am blood glucose levels were measured with a glucometer (Diathrive, South Salt Lake, Utah, USA) by obtaining a drop of blood from the tail. Subsequently, mice received 2 g of glucose (Sigma G8270) / kg of body weight by intraperitoneal injection. Blood glucose levels were measured again after 15, 30, 60, and 120 min after injection.

To measure insulin levels 20 µl of blood were collected from the tail before glucose injection and on every following time point and mixed with heparin. The blood plasma was prepared by centrifugation at 2000 x g for 15 minutes. The insulin concentration in blood plasma was determined using the Ultra Sensitive Mouse Insulin ELISA Kit (Cat. 90080, Crystal Chem, Illinois, USA) following the manufacturer’s instructions.

### Insulin Tolerance Test

Before the procedure mice were fasted for 6h in clean cages with no food or feces in the bedding starting at 9am. Blood glucose levels were measured with a glucometer (Diathrive, South Salt Lake, Utah, USA) by obtaining a drop of blood from the tail. Subsequently, mice were injected with 0.75 units/kg body weight of human insulin (Sigma I9278) by intraperitoneal injection. Blood glucose levels were measured again after 15, 30, 45, 60, and 120 min after injection.

To measure insulin levels 20 µl of blood were collected from the tail before glucose injection and on every following time point and mixed with heparin. The blood plasma was prepared by centrifugation at 2000 x g for 15 minutes. The insulin concentration in blood plasma was determined using the Ultra Sensitive Mouse Insulin ELISA Kit (Cat. 90080, Crystal Chem, Illinois, USA) following the manufacturer’s instructions.

### Measuring concentrations of FFA, glycerol, and triglycerides

FFA contents in blood serum were determined the Free Fatty Acid Quantitation Kit (Sigma, MAK044) following the manufacturers’ protocols. Glycerol and triglyceride concentrations in blood serum were measured using the Triglyceride Determination Kit (Sigma, TR0100) following the manufacturer’s instructions.

### Histology

For histological examination interscapular brown adipose tissue (BAT), inguinal white adipose tissue (sWAT), and epididymal white adipose tissue (vWAT) were fixed in 4% formalin for 24h directly after dissection. Tissues were rinsed and stored in 95% until embedding in paraffin. Paraffin blocks were sectioned (10µm) and sections mounted on glass slides and stained with hematoxylin and eosin (H & E). Images were taken at 20x magnification and the adipocyte area was calculated using the Fiji software plug-in Adiposoft (Galarraga et al., 2012).

### Reverse transcription quantitative polymerase chain reaction

Total RNA was extracted from fat tissues using the RNeasy Lipid Tissue Mini Kit (Qiagen, Hilden, Germany) and from liver using the RNeasy Mini Kit (Qiagen, Hilden, Germany). Equal amounts of RNA were reverse transcribed to cDNA using the iScript cDNA Synthesis system (Bio-Rad, Hercules, CA, USA). Relative transcript amounts were measured by qPCR using PowerUp SYBR Green Master Mix (Appliedbiosystems, Vilnius, Lithuania). All expression data were normalized to TATA-binding protein (Tbp) expression. The primer sequences were as follows: Cd36 forward → 5’-ATG GGC TGT GAT CGG AAC TG-3’; Cd36 reverse → 5’-TTT GCC ACG TCA TCT GGG TTT-3’; Tbp forward → 5’-GAA GCT GCG GTA CAA TTC CAG-3’; Tbp reverse → 5’-CCC CTT GTA CCC TTC ACC AAT-3’; Ucp1 forward → 5’-AGC CGG CTT AAT GAC TGG AG-3’; Ucp1 reverse → 5’-TCT GTA GGC TGC CCA ATG AAC-3’; Angptl4 forward → 5’-CAT CCT GGG ACG AGA TGA ACT -3’; Angptl4 reverse → 5’-TGA CAA GCG TTA CCA CAG GC -3’; Pepck forward → 5’-CTG CAT AAC GGT CTG GAC TTC -3’; Pepck reverse → 5’-GCC TTC CAC GAA CTT CCT CAC -3’; Lipe (Hsl) forward → 5’-GAT TTA CGC ACG ATG ACA CAG T -3’; Lipe (Hsl) reverse → 5’-ACC TGC AAA GAC ATT AGA CAG C -3’; Glut4 forward → 5’-ATC ATC CGG AAC CTG GAG G -3’; Glut4 reverse → 5’-CGG TCA GGC GCT TTA GAC TC -3’; Srebp1 forward → 5’-TGA CCC GGC TAT TCC GTG A -3’; Srebp1 reverse → 5’-CTG GGC TGA GCA ATA CAG TTC -3’; Fasn forward → 5’-GGA GGT GGT GAT AGC CGG TAT -3’; Fasn reverse → 5’-TGG GTA ATC CAT AGA GCC CAG -3’; Hsd11ß1 forward → 5’-GGA GCC CAT GTG GTA TTG ACT -3’; Hsd11ß1 reverse → 5’-CCG CAA ATG TCA TGT CTT CCA T -3’; Dgat2 forward → 5’-GCG CTA CTT CCG AGA CTA CTT -3’; Dgat2 reverse → 5’-GGG CCT TAT GCC AGG AAA CT -3’; Atgl (Pnpla2) forward → 5’-ATG TTC CCG AGG GAG ACC AA -3’; Atgl (Pnpla2) reverse → 5’-GAG GCT CCG TAG ATG TGA GTG -3’.

### RNAseq

RNA was extracted from BAT or vWAT using the RNeasy Lipid Tissue Mini Kit (Qiagen, Hilden, Germany). For each condition (sham pellet, corticosterone pellet, sham injection, corticosterone injection) RNA was extracted from 4 mice per time point (3, 7, 14 days). RNA was also isolated from 4 mice before start of treatment (day 0). RNA quality of all samples was evaluated by Bioanalyzer 2100 High Sensitivity RNA Analysis chips (Agilent, Cat. 5067-1513) which displayed intact RNA integrity. mRNA samples were concentrated to ≤ 5 µl by MinElute column (QIAGEN, Cat. 74204). For generation of RNA-seq libraries, polyadenylated mRNA was isolated from 300 ng of total RNA by incubation with oligo-DT attached magnetic beads and followed by strand-specific library preparation using the TruSeq Stranded mRNA Library Preparation kit (Ilumina, Cat. 20020595). Briefly, isolated polyadenylated mRNA was fragmented using divalent cations under elevated temperature and 1st and 2nd strands DNA were synthesized using SuperScript II Reverse Transcriptase (provided with Ilumina kit). A-tailing and adapter ligation was performed according to the manufacturer’s protocol; the resulting dsDNA was enriched in a PCR reaction based on predetermined CT values and cleaned using AMPure XP beads (provided with Ilumina kit). Concentrations of enriched dsDNA fragments with specific adapters were determined and base pair average size and library integrity were analyzed using the Bioanalyzer DNA High Sensitivity chips (Agilent, Cat. 5067-4626). Samples were pooled and sequenced on the Illumina NextSeq 500/550 High Output platform (Illumina, FC-404-2002) up to 18 samples per lane with 1% PhiX spike as a control.

To assess the read quality of the raw FASTQ files we used FastQC (Andrews and Babraham Bioinformatics, 2010) (v0.11.7). Pseudo-alignment of the reads to the mouse reference transcriptome (Mus_musculus.GRCm38.cdna) was carried out using Kallisto (Bray et al., 2016) (v0.44.0) with the quantification algorithm enabled, the number of bootstraps set to 100, and ran in paired-end mode. The Kallisto output files imported to R with Sleuth, and the transcripts per million (TPM) were used for downstream differential expression analysis (Pimentel et al., 2017).

We used the Sleuth package to identify differentially expressed genes with the full model consisting of treatment, lane, replicate, and day and the reduced model consisting of lane and replicate. This allowed us to identify genes that were differentially expressed based on treatment (i.e. corticosterone pellet vs sham pellet) and time. All downstream analysis was done on transcripts that were considered significant from the DESeq2 analysis (likelihood ratio test with a FDR less than 0.05). Outlier samples were identified by both clustering and PCA analysis and removed. The number of differentially expressed genes after removing outliers increased by 1,668, with all but 31 genes common between the two models.

For more information, please see https://kylekovary.shinyapps.io/Flattened-diurnal-glucocorticoid-oscillations/.

### Gene Ontology (GO) Analysis

GO term analysis was done using the clusterProfiler package(Yu et al., 2012). GO terms of interest were identified by calculating the adjusted p-value between two sample groups (i.e. corticosterone pellet vs. sham pellet) as well as a z score for general trends of increased or decreased differential expression of the genes within the GO terms. The z score is defined as the number of genes within the GO term with an absolute value log2 fold change greater than or equal to 1.5 (fold change > 1.5 or < 0.67) divided by the square root of the total number of genes assigned to that GO term(Walter et al., 2015).

### Western Blot Analysis

For western blot analysis 10mg of fat tissue were lysed in 50 mM Tris-HCl, 1 % Nonidet P-40, 0.1 % sodium dodecyl sulfate (SDS) and 0.15 M NaCl, pH 8, supplemented with protease inhibitors (cOmplete, Mini, EDTA-free, Sigma Aldrich). For preparation of tissue lysates, a homogenizer (Fisher Scientific) was used and cell debris removed by centrifugation at max speed for 20 min at 4 °C. Protein concentrations were determined using the bicinchoninic acid (BCA) protein assay kit (Pierce, Rockford, USA). 50 μg of proteins of tissue lysates were boiled for 5 min at 95°C in NuPAGE LDS sample buffer containing 1x reducing agent (Invitrogen). Lysates were separated by SDS–PAGE using the XCell SureLock Electrophoresis Cell on NuPAGE Novex 4%–12% Bis-Tris Protein Gels (Invitrogen). Proteins were transferred onto polyvinylidene fluoride membranes (Thermo Scientific) using the XCell II Blot Module (Invitrogen). Membranes were blocked by incubating with TBS with 0.1% Tween-20 containing 3% non-fat milk for 2h at room temperature or overnight at 4°C. After blocking, membranes were exposed to the primary antibody (UCP-1; used 1:800) overnight at 4°C. After washing (TBS with 0.1% Tween-20), the membranes were incubated for 2 h with the secondary antibody. The membranes were washed (TBS with 0.1% Tween-20) and developed with the Supersignal West Femto chemiluminescence substrate (Pierce, Rockford, USA). Peroxidase activity was detected with a Lumi Imager device (ImageQuant LAS 4000mini, GE Healthcare). The primary antibody UCP-1 was purchased from α-Diagnostics (Catalog No. UCP11-A). The secondary antibody was purchased from Cell Signaling (α-rabbit: Catalog No. #7074).

### Statistics

All data are represented as mean ± SD or mean ± SEM and analyzed by Students t test using GraphPad Prism software. n indicates the number of animals per group or number of independent experiments. Results were considered significant if p < 0.05.

## Notes

### Competing Interest Statement

The authors have declared no competing interest.

