## Supplementary material for "Flattening of diurnal glucocorticoid oscillations causes Cd36 and insulin-mediated obesity": Supp Table 1

| | Wild-type mice | | | | Cd36ko mice | |
|---|---|---|---|---|---|---|
| | **Sham pellet** | **Cort pellet** | **Sham injection** | **Cort injection** | **Sham pellet** | **Cort pellet** |
| **Body weight gain [g]** | 2.7 ± 0.5 | 4.8 ± 0.5 | 1.0 ± 0.3 | 1.1 ± 0.3 | 1.7 ± 0.4 | 3.3 ± 0.4 |
| **Body weight gain [% of initial body weight]** | 11.0 ± 0.8 | 20.1 ± 1.5 | 4.4 ± 1.4 | 4.4 ± 1.0 | 7.8 ± 1.0 | 14.7 ± 1.9 |
| **sWAT mass [mg]** | 296.2 ± 27.4 | 721.2 ± 35.1 | 326.0 ± 72.5 | 329.0 ± 6.4 | 264.0 ± 23.6 | 557.3 ± 49.9 |
| **vWAT mass [mg]** | 448.8 ± 37.6 | 1250.0 ± 45.5 | 456.3 ± 105.7 | 496.0 ± 27.8 | 468.2 ± 60.3 | 676.3 ± 48.8 |
| **BAT mass [mg]** | 148.8 ± 12.5 | 321.4 ± 11.5 | 123.7 ± 16.6 | 151.0 ± 10.6 | 98.0 ± 10.2 | 178.3 ± 12.6 |
| **Insulin [ng/ml]; fasted** | 0.52 ± 0.06 | 1.53 ± 0.16 | 0.56 ± 0.08 | 0.54 ± 0.04 | n.d. | n.d. |
| **Insulin [ng/ml]; not fasted** | 0.61 ± 0.06 | 2.60 ± 0.72 | n.d. | n.d. | 1.52 ± 0.22 | 7.43 ± 1.35 |
| **Blood glucose [mg/dl]; fasted** | 131.0 ± 22.1 | 139.8 ± 24.0 | 161.0 ± 15.1 | 160.7 ± 14.7 | n.d. | n.d. |
| **Blood glucose [mg/dl]; not fasted** | 189.1 ± 11.0 | 146.3 ± 8.3 | n.d. | n.d. | 171.5 ± 8.4 | 140.0 ± 5.0 |
| **Liver [mg]** | 971.3 ± 48.5 | 974.0 ± 9.5 | 977.0 ± 51.4 | 1027.3 ± 38.5 | n.d. | n.d. |
| **Food intake [g/mouse/day]** | 3.5 ± 0.1 | 3.8 ± 0.1 | 3.1 ± 0.1 | 3.2 ± 0.1 | n.d. | n.d. |
| **FFA [mM]** | 0.84 ± 0.13 | 1.65 ± 0.08 | 1.21 ± 0.24 | 1.16 ± 0.15 | n.d. | n.d. |
| **Glycerol [mg/ml]** | 0.29 ± 0.05 | 0.48 ± 0.05 | 0.35 ± 0.03 | 0.46 ± 0.04 | n.d. | n.d. |
| **Triglycerides [mg/dl]** | 46.2 ± 6.8 | 72.6 ± 6.0 | 84.8 ± 6.5 | 79.1 ± 7.6 | n.d. | n.d. |

**Table S1: Summary of mouse experiments.** All data was acquired after a treatment period of 21 days, except data for blood glucose (not fasted), FFA, glycerol, and triglyceride levels, which were determined 14 days after beginning of treatment. Values are presented as mean +/- SEM; n.d. = no data.
