## Supplemental Figures for "Flattening of diurnal glucocorticoid oscillations causes Cd36 and insulin-mediated obesity"

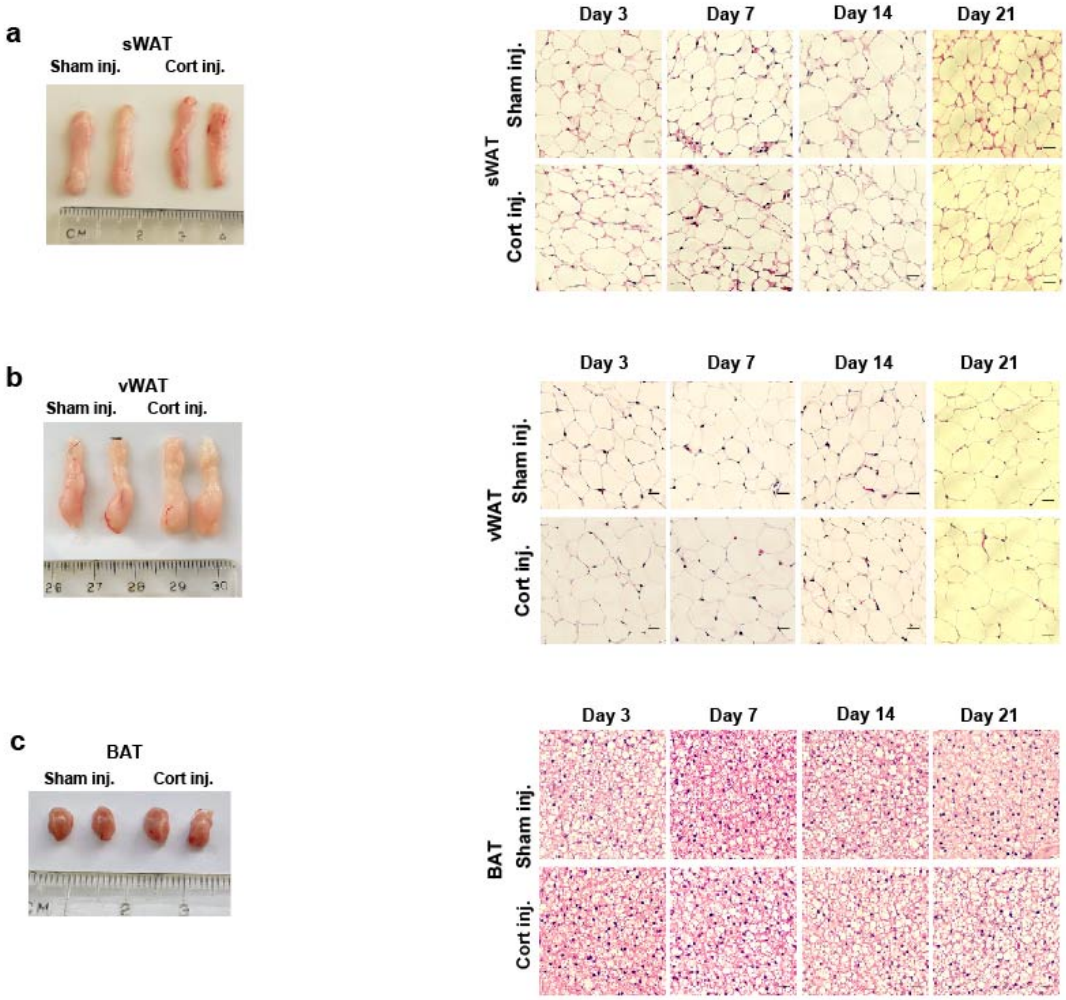


**Figure S1. Images of tissue and adipocyte sections in WAT and BAT induced by daily corticosterone injection versus sham injection.**

(A-C) Lack of a visible change in size of WAT and BAT tissue and size of adipocytes in Cort injected versus Sham injected mice (n=3 mice per treatment, 25 images per mouse (20x)).

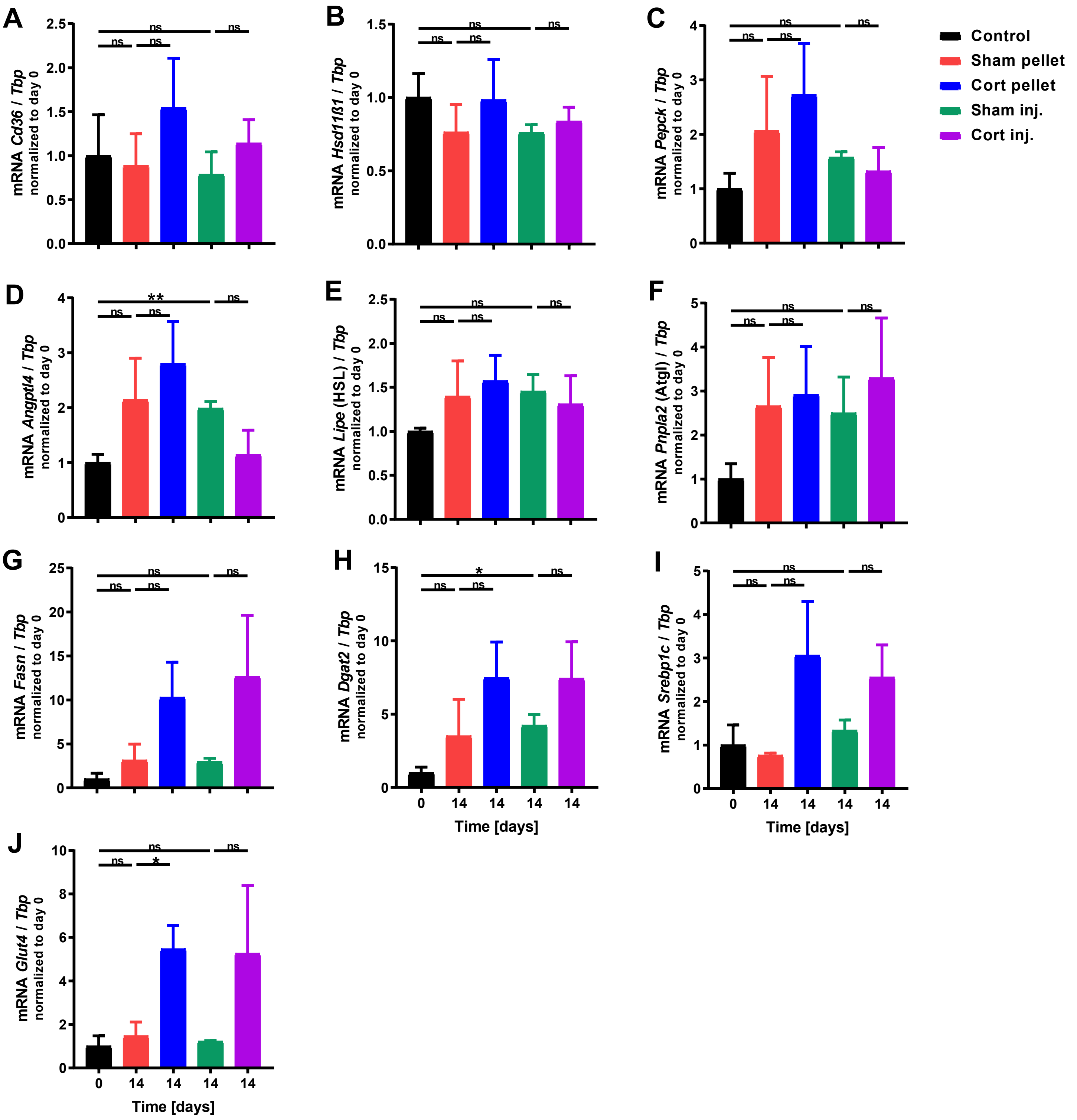


**Figure S2. Flattening of corticosterone oscillations does not result in significant gene expression alterations of genes involved in lipid metabolism in liver.**

(A-J) Gene expression of different genes measured by qPCR in liver after flattening corticosterone levels for 14 days. Gene expression data was normalized to the expression of Tbp and is presented as mean ± SEM (n = 3-4). Unpaired t test, $*p < 0.05$, $**p < 0.01$, ns = not significant. Control group are mice at beginning of experiment (day 0).

**a**

| depot | cond_1 | cond_2 | num_sig |
|---|---|---|---|
| BAT | Cort injection | Cort pellet | 2866 |
| BAT | Cort injection | Sham injection | 9 |
| BAT | Cort injection | Sham pellet | 110 |
| BAT | Cort pellet | Sham pellet | 7146 |
| BAT | Sham injection | Cort pellet | 3502 |
| BAT | Sham injection | Sham pellet | 2 |
| vWAT | Cort injection | Cort pellet | 4022 |
| vWAT | Cort injection | Sham injection | 0 |
| vWAT | Cort injection | Sham pellet | 0 |
| vWAT | Cort pellet | Sham pellet | 6233 |
| vWAT | Sham injection | Cort pellet | 9800 |
| vWAT | Sham injection | Sham pellet | 0 |

**b**

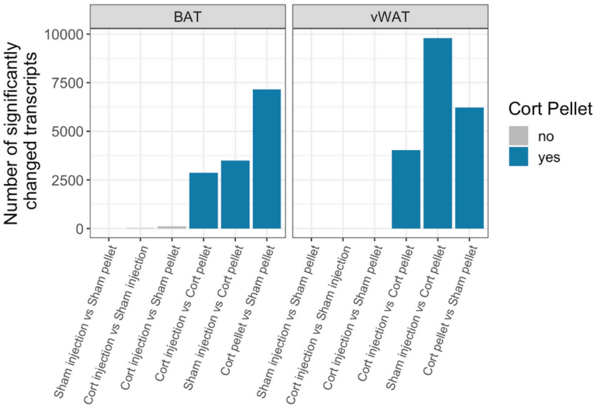


As you can see, in both tissues the main driver of differential expression of mRNA is the cort pellet. Since, in our opinion, the placebo pellet is the best control condition to compare the cort pellet against, it is what we will be comparing the cort pellet against in WAT and BAT tissue going forward.

**c**

Overlap between the up regulated genes between WAT and BAT

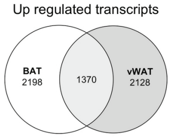


Overlap between the down regulated genes between WAT and BAT

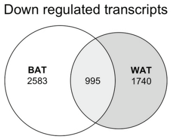


**Figure S3. Comparison of BAT and vWAT RNAseq datasets.**
(A-B) Number of significantly changed transcripts in BAT and vWAT in different comparisons of treatment methods.
(C) Number of overlapping transcripts in vWAT and BAT. See also Tables S2-S7.

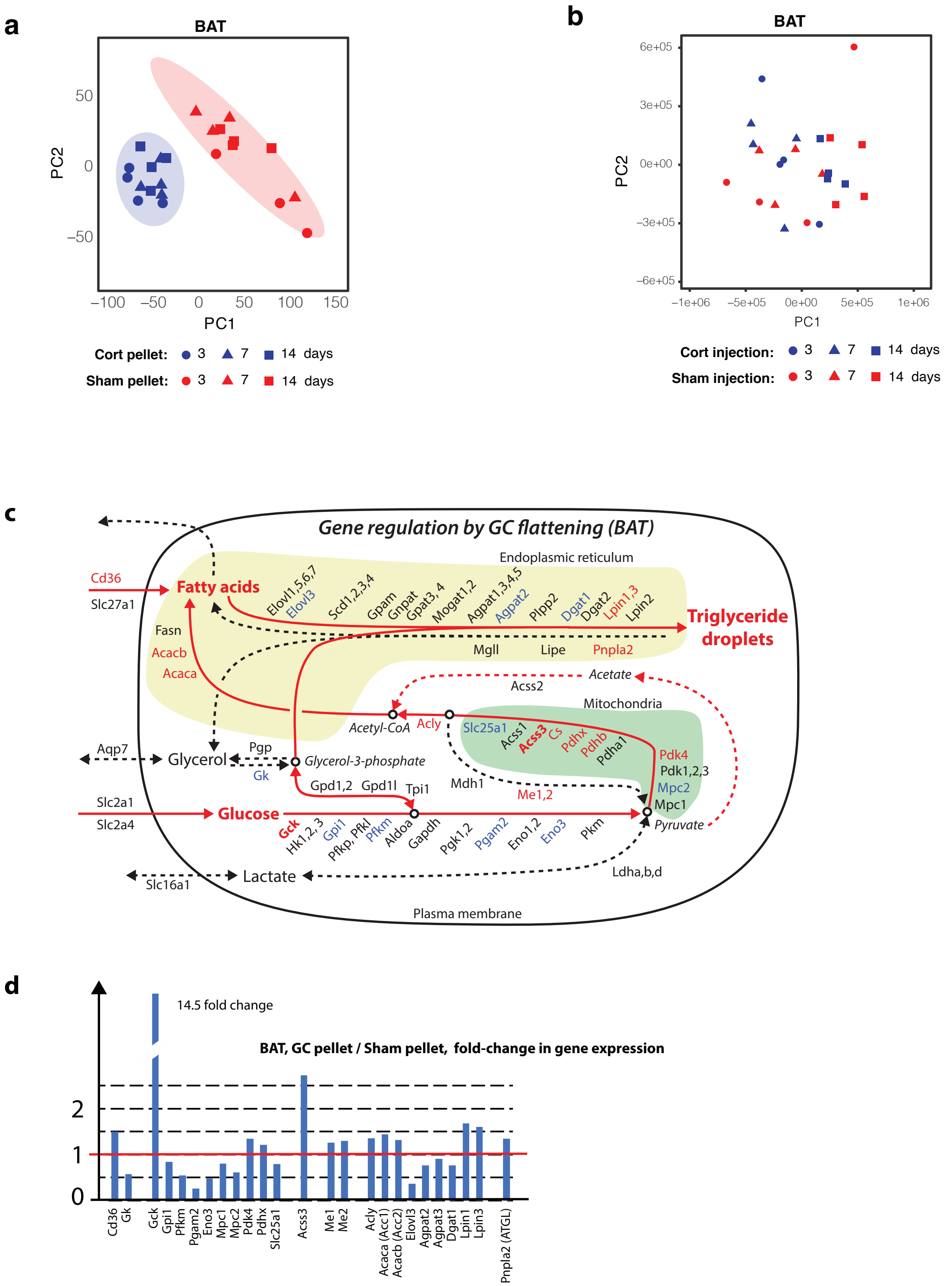


**Figure S4. Analysis of BAT RNAseq dataset.**

(A) Principal component analysis of BAT from corticosterone versus sham pellet-implanted mice obtained at 3, 7, and 14 days after pellet implantation.

(B) Principal component analysis of BAT from corticosterone and sham injected mice obtained at 3, 7, and 14 days after the start of injections.

(C) Schematic of gene expression changes in triglyceride synthesis pathways in BAT from corticosterone versus sham pellet-implanted mice. Upregulated, unchanged, and downregulated genes are marked in red, black, and blue, respectively. Bold type mark genes with a more than 2-fold change in gene expression. See also Tables S2-S7.

(D) Barplots showing fold-change of significantly changed BAT genes in corticosterone versus sham pellet-implanted mice. See also Tables S2-S7.

**a**

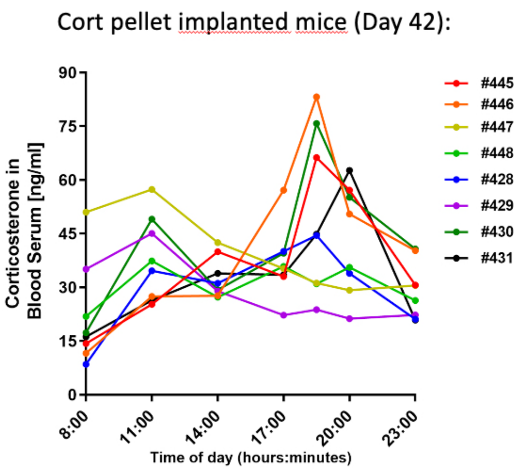


**b**

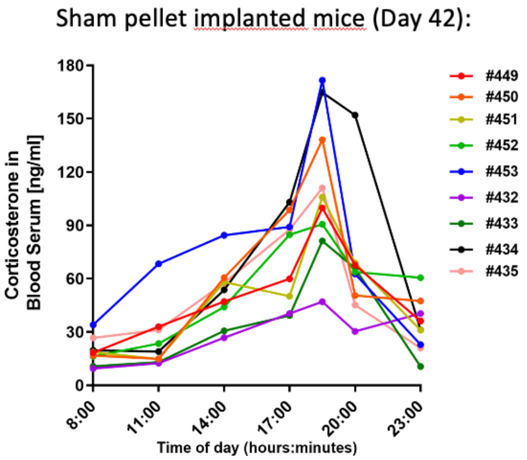


**Figure S5. Corticosterone measurements in mice implanted with corticosterone or sham pellets at Day 42 after pellet implantation.**

(A) Blood serum corticosterone levels at Day 42 for 8 corticosterone pellet-implanted mice.

(B) Blood serum corticosterone levels at Day 42 for 8 sham pellet-implanted mice.

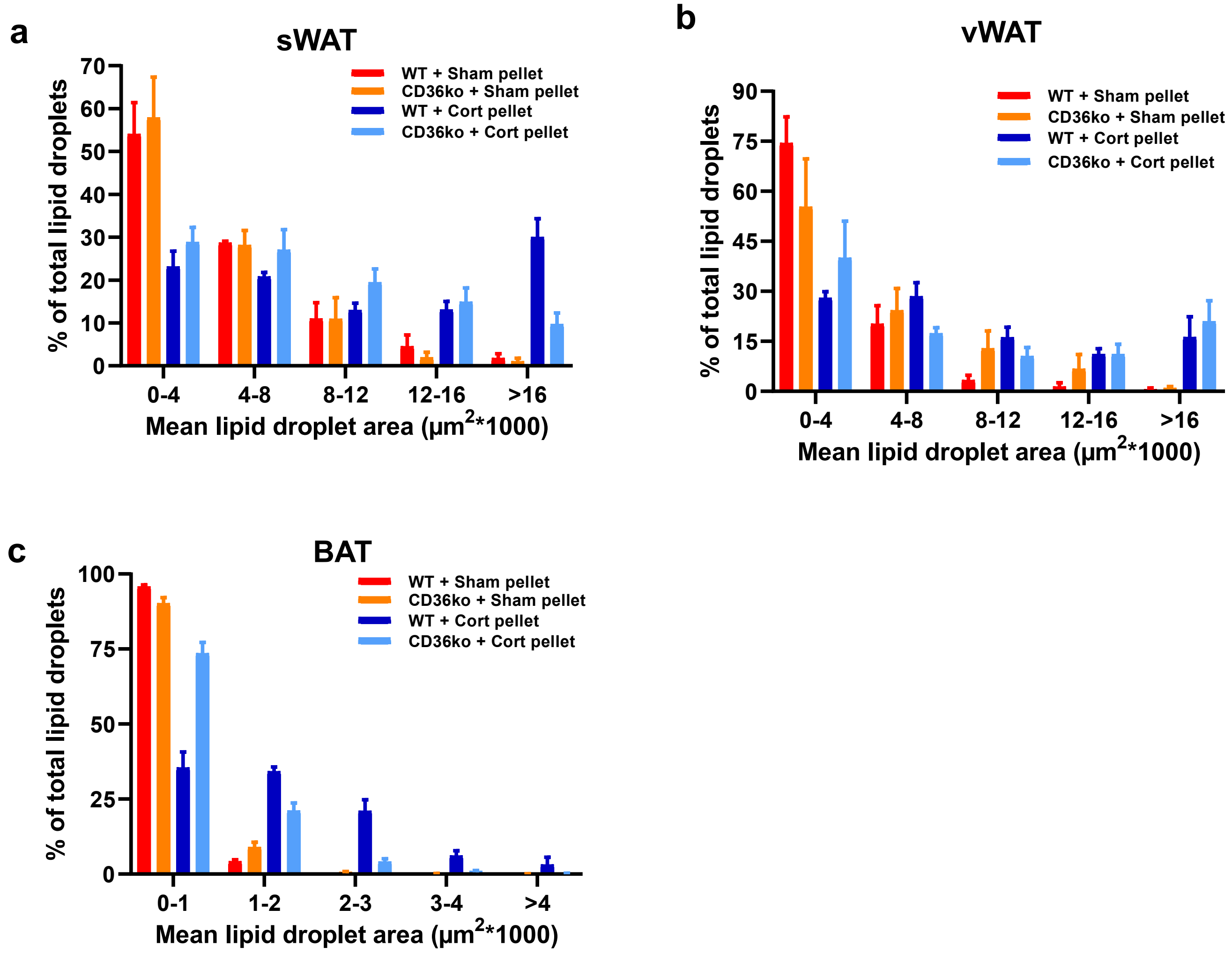


**Figure S6. Quantification of lipid droplet size in WT and CD36KO mice implanted with corticosterone or sham pellets for 21 days.**
(A-C) Frequency distributions of mean lipid droplet area in sWAT, vWAT, and BAT as analyzed using Adiposoft (mean + SEM; n=3 mice per treatment, 25 images per mouse (20x)).
.
